# Induced B-Cell Receptor Diversity Predicts PD-1 Blockade Immunotherapy Response

**DOI:** 10.1101/2024.12.03.626669

**Authors:** Yonglu Che, Jinwoo Lee, Farah Abou-Taleb, Kerri E. Rieger, Ansuman T. Satpathy, Anne Lynn S. Chang, Howard Y. Chang

## Abstract

Immune checkpoint inhibitors such as anti-PD-1 antibodies (aPD1) can be effective in treating advanced cancers. However, many patients do not respond and the mechanisms underlying these differences remain incompletely understood. In this study, we profile a cohort of patients with locally-advanced or metastatic basal cell carcinoma undergoing aPD-1 therapy using single-cell RNA sequencing, high-definition spatial transcriptomics in tumors and draining lymph nodes, and spatial immunoreceptor profiling, with long-term clinical follow-up. We find that successful responses to PD-1 inhibition are characterized by an induction of B-cell receptor (BCR) clonal diversity after treatment initiation. These induced BCR clones spatially co-localize with T-cell clones, facilitate their activation, and traffic alongside them between tumor and draining lymph nodes to enhance tumor clearance. Furthermore, we validated aPD1-induced BCR diversity as a predictor of clinical response in a larger cohort of glioblastoma, melanoma, and head and neck squamous cell carcinoma patients, suggesting that this is a generalizable predictor of treatment response across many types of cancers. We discover that pre-treatment tumors harbor a characteristic gene expression signature that portends a higher probability of inducing BCR clonal diversity after aPD-1 therapy, and we develop a machine learning model that predicts PD-1-induced BCR clonal diversity from baseline tumor RNA sequencing. These findings underscore a dynamic role of B cell diversity during immunotherapy, highlighting its importance as a prognostic marker and a potential target for intervention in non-responders.

## INTRODUCTION

Immune checkpoint inhibitors, including programmed cell death protein 1 (PD-1) inhibitors, have significantly improved long-term outcomes of patients with advanced cancer. Despite this success, patient responses are highly variable, and the underlying mechanisms of a successful response are incompletely understood (Li et al., 2022, Warner et al., 2020, Sharma et al., 2017). Many studies have attempted to identify predictors of treatment response, including tumor expression of PD-L1, PD-L2, IFNγ, CD73, and CXCL9, as well as tumor mutation burden and the formation of tertiary lymphoid structures, but these do not fully explain the variability of patient responses (Meliante et al., 2023, Cristescu et al., 2018, Sautès-Fridman et al., 2016).

The tumor microenvironment is highly dynamic. Contrary to the classical view of immunologically “hot” vs. “cold” tumors defined by the mere presence or absence of T cell infiltration, we and others discovered that aPD1 checkpoint blockade induced clonal replacement of T cells, subsequent expansion detectable in the peripheral blood, migration into the tumor, followed by multiple rounds of activation, expansion, and ultimately T cell exhaustion (Yost et al., 2019, Chaft et al., 2022). These dynamic changes highlight the importance of immune cell trafficking and nominate tumor draining lymph nodes as compelling sites of cell-cell interactions (Yost et al., 2021). Recent evidence suggests that B cells play a crucial role in the anti-PD-1 response. In melanoma patients, B cell markers are enriched in responders and localize to tumor tertiary lymphoid structures (Helmink et al., 2020). Similarly in soft tissue sarcomas, B-cell-rich tertiary lymphoid structures were associated with better prognosis (Petitprez et al., 2020, Carbrita et al., 2020). Additional studies have implicated specific B cell subtypes as positive prognostic indicators in multiple cancer types (Yang et al., 2024, Ding et al., 2023, Griss et al., 2019, Singh et al., 2022, Zhang et al., 2023, Shu et al., 2024, Gavrielatou et al., 2024, Chang et al., 2024), further highlighting the importance of B cells in antitumor immunity. Questions remain, however, regarding the mechanisms behind these findings and how these insights might translate into clinical practice.

Building on our previous longitudinal analysis, which examined the role of novel tumor-infiltrating T cell clonotypes in reinvigorating anti-tumor responses following PD-1 blockade in advanced cutaneous basal cell carcinoma (BCC) and cutaneous squamous cell carcinoma (SCC) (Yost et al., 2019), we performed long-term clinical outcome monitoring, single cell transcriptome and immunoreceptor sequencing, and high-definition spatial transcriptomics of tumors and draining lymph node specimens collected over years of direct patient care. Because T- and B-cell receptor genes undergo somatic recombination, TCR and BCR sequences provide unique molecular barcodes that are passed onto their progenies, allowing us to track the clonal dynamics during checkpoint blockade in patients. By integrating single-cell and spatial transcriptomic profiling with clinical outcomes, we reveal a significant role for increased B-cell receptor (BCR) clonal diversity following PD-1 blockade. Our findings demonstrate that diverse B-cell clones migrate and interact with specific T cells in both the tumoral microenvironment and within the draining lymph nodes to enhance T-cell expansion and activation, ultimately improving tumor clearance.

## RESULTS

### Patient Survival after PD-1 Inhibition in Basal Cell Carcinoma Is Associated with Intratumoral BCR Diversification

Previous work from our group led to the discovery that expansion of tumor-infiltrating T lymphocytes arises from novel rather than pre-existing clones in a cohort of locally advanced and metastatic cutaneous carcinoma patients treated with PD-1 inhibitors (Yost et al., 2019). To gain further insight into the mechanisms that influence long-term survival after treatment, we extended the clinical follow-up of this patient cohort and archived additional samples taken for clinical care during this time for future analysis (**Figure 1A, Supplemental Data Table 1**). To investigate B cell clonality within our cohort, we analyzed single-cell RNA sequencing data to identify BCR sequences and quantify the number of unique clonotypes per tumor specimen (**Figure 1B-C, Extended Data Figure 1 A-F**). The sequencing datasets were prepared using 5’ gene expression library construction as previously described and sequenced to a minimum depth of 25,000 reads per cell. Using TRUST4 for clonotype calling analogous to previous work in this field (Song et al., 2021, Helmink et al. 2020), we find that the complementarity-determining region 3 (CDR3) sequence lengths in our analysis displayed comparable distributions to known BCR and TCR lengths (DeWitt et al. 2016).

**Figure 1:**
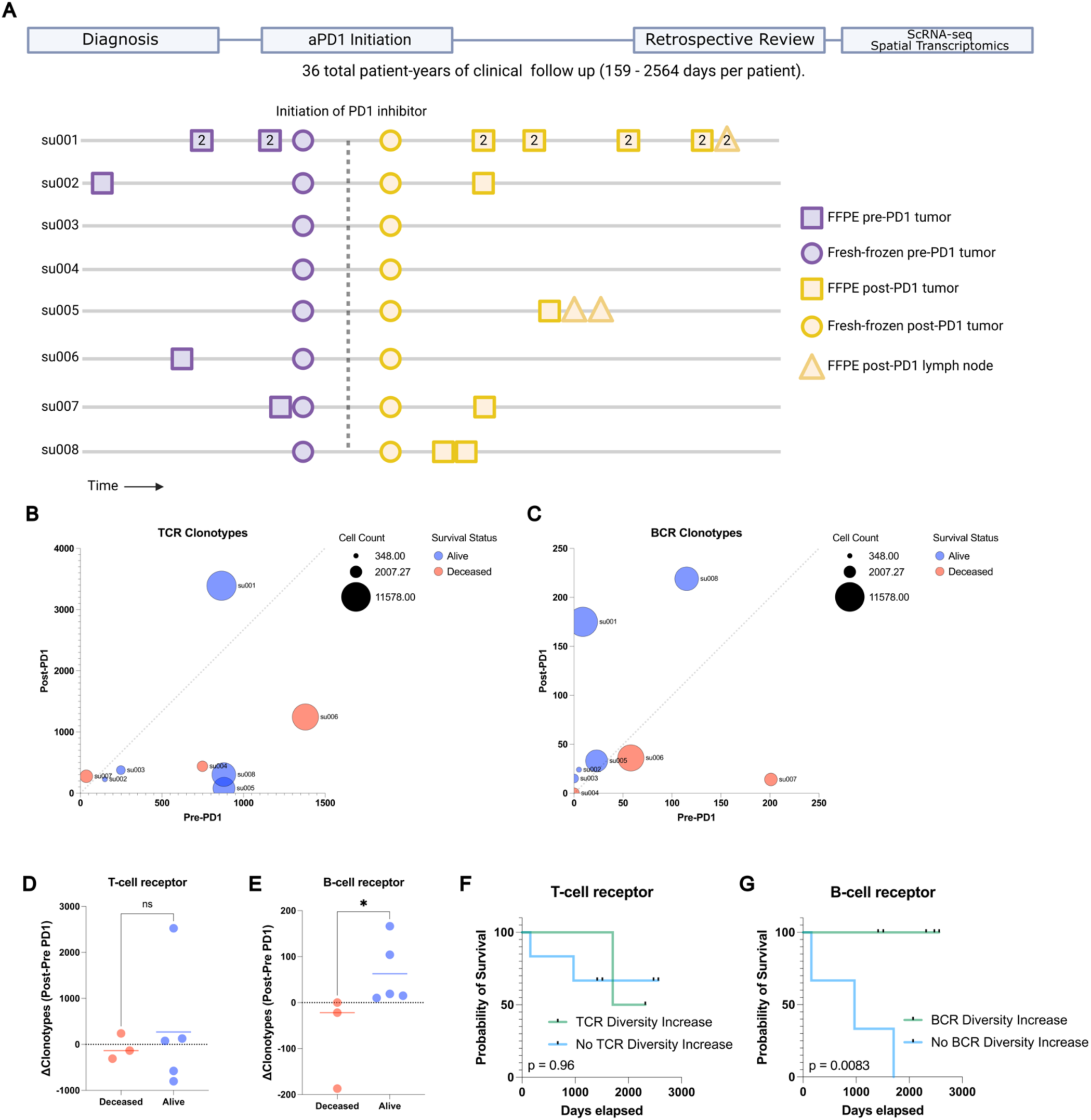
Long term survival of advanced BCC patients highlights prognostic role of aPD1-induced BCR clonal diversity. A) A schematic diagram of our BCC patient cohort. Top: A retrospective chart review was performed after an extended period of clinical follow up lasting up to 2564 days to allow final clinical outcomes of PD-1 blockade to manifest. Bottom: A visualization of the samples taken for our prior study (Yost et al. 2019) and this study. All patients had pre- and post-PD-1 inhibitor-treated tumor samples for scRNA-seq (circles). All available archived specimens in long-term storage were accessed (squares and triangles) and processed for spatial transcriptomics. Numbers within the shapes represent replicates taken from sequential sections of the sample tissue block. Y axis represents anonymized patient identifiers while the x-axis represents time (not to exact scale). B) Unique TCR and C) BCR clonotype counts detected per patient in their pre-PD-1 inhibitor tumor (X-axis) and post-PD-1 inhibitor treated tumor (Y axis) as calculated by TRUST4 run on 5’ single cell RNA sequencing as described (Methods). Point sizes represent the total cell count obtained from tumors of each patient. D) Change in unique TCR and E) BCR clonotype counts between pre-PD-1 inhibitor and post-PD-1 inhibitor tumors for each patient (Y-axis), stratified by clinical status at last available follow up. F) Kaplan-Meier curve of overall survival for our BCC patient cohort stratified by the change in pre-to-post PD-1 inhibitor TCR and G) BCR clonotype count. P-values calculated using the log-rank test.

Analysis of the TCR and BCR repertoires revealed considerable heterogeneity both among patients and between pre- and post-PD-1 inhibitor treatment tumor specimens from the same patient (**Figure 1 B-C**), suggesting a level of stochasticity in the tumor-infiltrating lymphocyte population. For clarity, we refer to “clonal expansion” or “clone size” as the number of T or B cells with the same immunoreceptor sequences, while “clonal diversity” refers to the number of T or B cells with distinct immunoreceptor sequences. TCR clonal expansion and TCR diversity were not directly associated with improved survival in this cohort (**Figure 1D**, **Extended Data Figure 1 G**). In contrast, while BCR clonotype counts at individual time points did not correlate with survival outcomes, we observed an increase in unique BCR clonotype counts after PD-1 blockade exclusively in surviving patients (**Figure 1 E**). Patients with an increase in intratumoral BCR clonotype diversity demonstrated a markedly improved survival rate compared to those without expansion (log-rank test p=0.0083, **Figure 1 F-G, Extended Data Figure 1H**), which contrasts with the lack of clear association between either TCR clonal expansion or TCR clonal diversity with survival outcomes. This led us to further investigate the mechanisms by which *PD-1 inhibition induces BCR diversity*.

### Targeted Spatial Transcriptomics of Archived Patient Specimens Before and After PD-1 Blockade Captures Evolution of the Tumor and Draining Lymph Node Microenvironment

To further explore mechanisms by which BCR clonotype diversity might enhance patient survival, we expanded our molecular profiling to include spatial transcriptomics on available specimens. Prior studies have highlighted the utility of spatial data to elucidate T/B cell anti-tumor mechanisms (Zhang et al. 2023), so we reasoned that this type of dataset would complement our existing scRNA-seq data. We selected the Xenium in-situ platform due to its 1) compatibility with formalin-fixed, paraffin-embedded (FFPE) archival specimens, 2) accurate cell segmentation, and 3) flexible custom probe design, which were critical features for our study (Janesick et al., 2023). Recognizing the platform’s current limitations in gene coverage, we carefully designed a custom probe panel to minimize information loss associated with this constraint (Methods). This was accomplished by constructing a 480 gene custom panel comprising the most differentially expressed genes in our scRNA sequencing, essential markers of cell type and cell state, and probes targeting TCR and BCR loci (detailed further below, Methods, **Extended Data Figure 2A, Supplemental Data Table 2**). To objectively evaluate this manually curated design, we then separately trained a neural net model to evaluate the panel’s potential performance. We assigned the model to predict the annotated cell states of the scRNA sequencing data, then measured the change in model accuracy when constrained to limited gene panels including our custom design (**Extended Data Figure 2B**). Notably, the 480 gene custom panel showed (under ideal, simulated circumstances) minimal information loss even when compared to the baseline model with full gene expression access and outperformed predesigned panel designs for our application including the recently released Xenium 5K gene panel.

Our final spatial transcriptomics dataset encompassed 35 individual tissue specimens including 29 tumor specimens (21 samples from 6 aPD1-treated patients, 8 samples from 4 aPD1-naïve patients), 2 specimens from a tumor-adjacent immune-related adverse event (irAE) (from a single patient), and 4 draining lymph node specimens (from 2 aPD1-treated patients), amounting to a total of 1,633,794 spatially resolved individual cells (**Extended Data Figure 3**). Combined with the scRNA sequencing data of 53,030 cells from the same patient cohort, this dataset serves as a high-depth resource for investigating tumor and lymph node evolution during checkpoint blockade. To demonstrate a method for integrated analysis of these two datasets, we employed ENVI--a computational tool designed for multimodal data integration of scRNA sequencing and spatial transcriptomics--to create a shared latent space between spatial and single-cell data (Haviv et al., 2024). This approach demonstrates that cells from both modalities can be merged for simultaneous analysis (**Figure 2A, Extended Data Figure 4 A-E**), facilitating a more detailed understanding of the tumor microenvironment and lymphatics.

**Figure 2:**
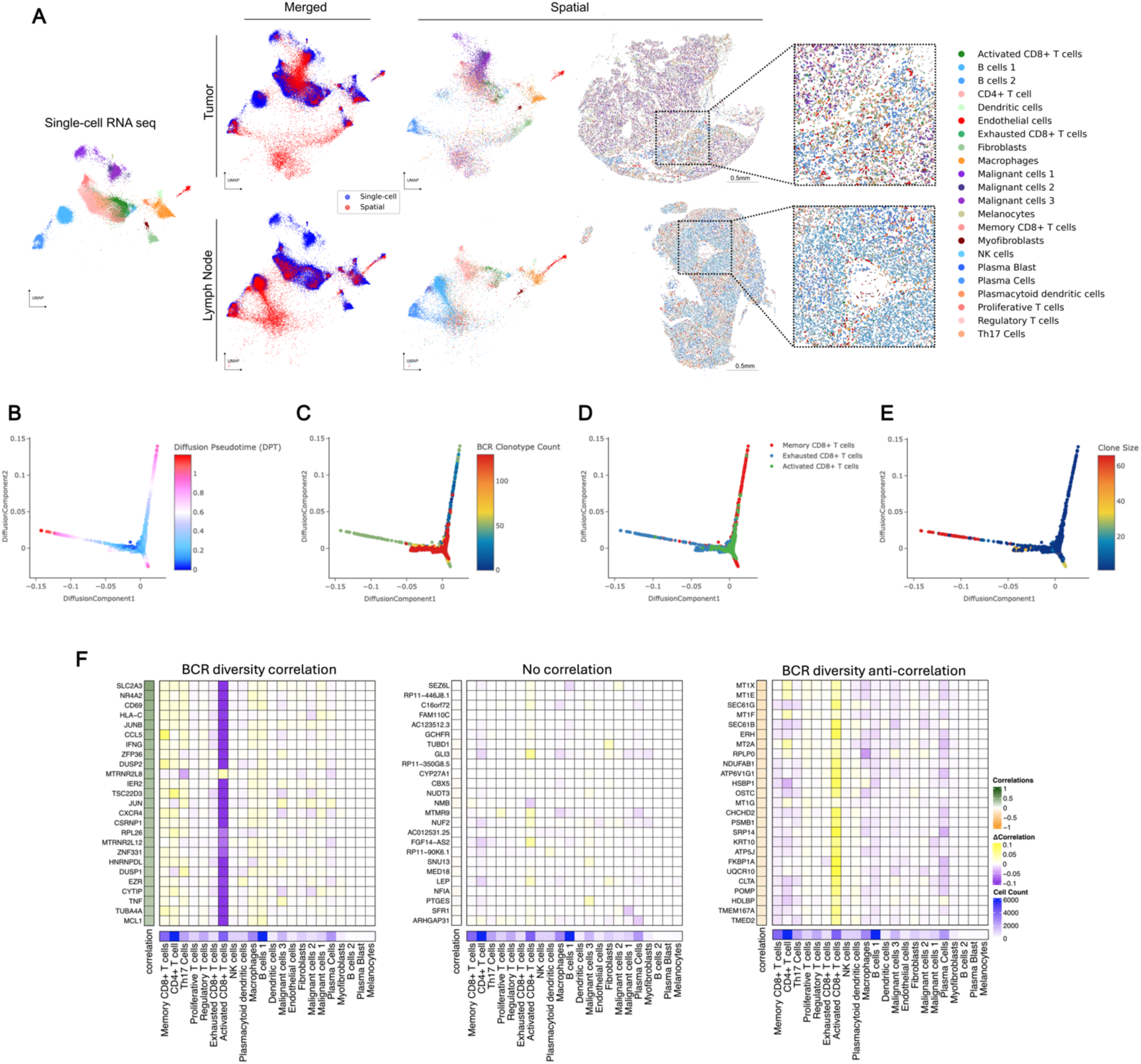
Spatial transcriptomics merged with scRNA-seq enables exploration of the BCR diversity-impacted tumor microenvironment. A) Representative tissue samples obtained from Xenium in-situ for patient su001 for tumor (top) and regional lymph node (bottom). Cells from Xenium were merged onto the same latent space (middle) as cells from our prior scRNA-seq (left and middle) using the computational tool ENVI (methods). Proposed cell types for Xenium cells were determined using a nearest-neighbor approach in the high-dimensional ENVI latent space. Xenium tissues are downsampled to 40% of total cells for visualization purposes. B) Diffusion plots of CD8+ T cell populations from scRNA-seq highlighting pseudotime, C) total unique BCR clonotype count present in the originating tumor of each T cell, D) annotated subtype of each T cell (bottom), E) and size of the originating T cell clonotype. F) Regression analysis of cell-type contributions to gene expression correlations to BCR-clonotype counts. Left heatmap represents the correlation across scRNA-seq tissue specimens between the annotated gene and the unique BCR clonotype count in that tissue specimen. The 25 most correlated genes (left), 25 most anti-correlated genes (right), and 25 random genes (middle) are shown. The middle heatmap grid represents the change in correlation strength of each gene (row) when a particular cluster (column) is blinded to the correlation calculation. The heatmap on the bottom row represents the total cell counts assigned to each cluster to verify that effect sizes are not solely influenced by cluster size.

### PD-1-induced B-Cell Diversity Supports Activation of CD8+ T cells

To investigate mechanisms by which B cells might influence tumor behavior, we examined relationships between BCR clonotype diversity and tumor composition using our scRNA-seq data. We isolated keratinocyte-containing cell clusters – which include the malignant basal cell carcinoma cells - and assessed their differentiation trajectories using diffusion maps (Haghverdi et al., 2015). BCC cells, characterized by stem-like properties resembling progenitor keratinocytes and expressing markers such as epithelial cell adhesion molecule EPCAM/BerEP4, mapped to earlier stages of differentiation in the diffusion plot (**Extended Data Figure 5A**). In contrast, terminally differentiating keratinocytes, likely from adjacent normal tissue captured in the samples, mapped to regions expressing higher levels of differentiation marker genes like keratin 1 (KRT1) and involucrin (IVL). Keratinocytes from tissue samples with higher BCR clonotype diversity were skewed towards terminal differentiation compared to those from low BCR clonotype diversity samples, suggesting lower tumor cell presence in these samples despite being biopsied from the center of clinically active disease.

We performed a similar analysis on B cells to assess the relationship between clonotype diversity and their differentiation states. Interestingly, samples with high BCR clonotype counts were skewed to antigen-presenting phenotypes rather than high immunoglobulin expression (**Extended Data Figure 5B**). These results support the hypothesis that diverse B cell clones are involved in antigen cross presentation to T cells.

Finally, we examined CD8+ T cells within the diffusion map overlaid with BCR clonotype diversity data. Our previously annotated subcategories of CD8+ T cell clusters--including activated, exhausted, and memory phenotypes--mapped to distinct differentiation trajectories. When overlaid with BCR clonotype counts of the originating samples, this analysis showed strong enrichment of the activated CD8+ T-cell phenotype in tumor samples with high BCR clonotype counts, suggesting that B-cell clonal diversity and CD8+ T-cell activation are closely linked (**Figure 2B-D, Extended Data Figure 5C**). When overlaid with T-cell clonotype sizes, we find that the clones with the largest sizes belonged to the exhausted CD8+ T-cell phenotype, whereas the clone sizes of activated CD8+ T cells tended to be smaller (**Figure 2E**). We pose that these T cells may be in the process of activation, supported by their diverse B cell environment, while exhausted T cells may be a signature more indicative of prior anti-tumor activity.

Using established network analysis approaches on our gene expression data, we identify pathways by which clonally diverse B cells can communicate with activated CD8+ T cells, including the major histocompatibility complex class I (MHC-I) pathway (**Extended Data Figure 6 A-D**), as well as validate the activation of the TCR signaling and activator protein 1 (AP-1) pathways in the activated CD8+ T-cell population (**Extended Data Figure 6 E-F**). To examine the specificity of the relationship between B cells and CD8+ T-cell activation, we performed a cluster-based regression analysis. We assessed correlations between gene expression in the scRNA sequencing dataset and the BCR clonotype counts of the originating samples (**Figure 2F**). Our aim was to determine whether gene expression patterns most correlated with intratumoral BCR clonotype diversity were attributable to a single cell type or influenced by cumulative shifts in multiple cell types. This analysis revealed a striking specificity originating from activated CD8+ T cells. The gene expression module correlating to high tumoral BCR diversity prominently featured the hallmark genes of T-cell activation such as CD69 (R= 0.42, FDR < 0.001), IFNG (R= 0.37, FDR < 0.001), and TNF (R= 0.29, FDR < 0.001), reinforcing the potential role of activated CD8+ T cells in mediating the effects of increased BCR diversity within the tumor microenvironment.

### Spatial Transcriptomics Enables Clonotype Tracking to Elucidate T-Cell:B-Cell Interactions

The strong correlation between BCR clonotype diversity and gene signature of activated CD8+ T cells, along with lower malignant keratinocyte burden in these BCR-diverse tissues, suggests a mechanism where B-cell clonal diversification following PD-1 blockade activates T cells leading to tumor clearance. Further exploration of this mechanism required analysis using spatial transcriptomics. This posed a technical challenge since established spatial technologies for archival FFPE specimens, to our knowledge, do not have the ability to simultaneously map lymphocyte clonotype locations. To overcome this, we further customized our Xenium panel to enable tracking of T and B cell clonotype trends in our tissue specimens.

While mapping specific CDR3 sequences is beyond the current limitations of the platform, we reasoned that variable (V)-gene usage from V(D)J-recombination events would vary between samples in a manner reflecting differences in lymphocyte clonotype composition. To validate this approach, we compared patient-matched and time-matched samples between the Xenium data and their corresponding scRNA-seq counterparts. Tumor samples from the same patient taken at the same clinical timepoint – pre- or post-PD-1 blockade – should show overlapping patterns of V-gene usage in the scRNA sequencing vs. Xenium in situ analyses. Indeed, this was true for both TCR alpha and TCR beta chains, where mismatched samples had no correlation in overlapping usage and matched samples demonstrated enrichment in shared V-gene expression patterns (**Figure 3A**). Interestingly, for BCRs, only the immunoglobulin (IG) heavy chain and not the IG light chain displayed this matched pattern (**Figure 3B**). Therefore, we proceeded with using the IG heavy chain alone for BCR clonotype tracking.

**Figure 3:**
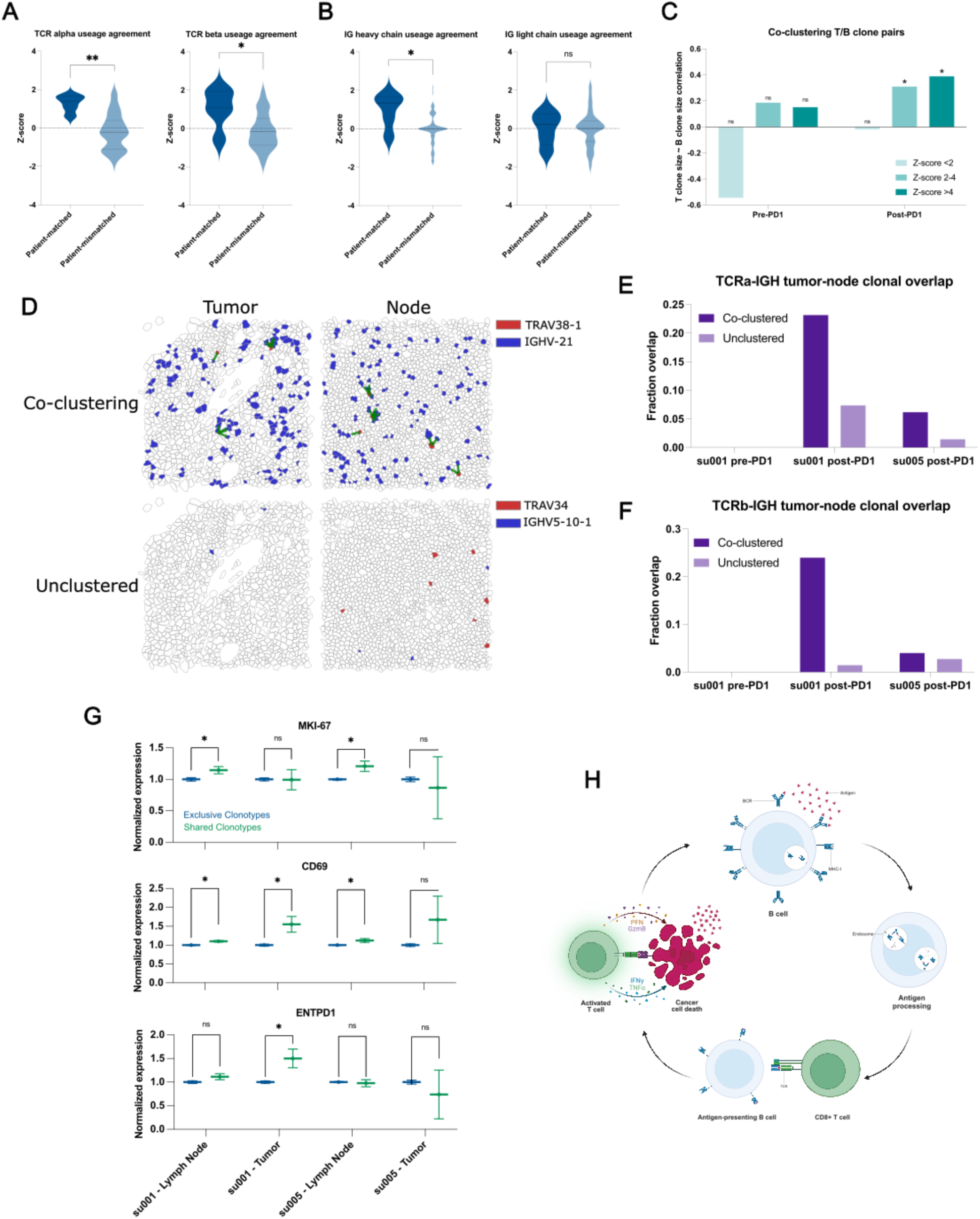
Clonally diverse B cells co-cluster with paired T cell clones with an activated phenotype. A) Correlations of V-gene usage between those detected using TRUST4 analysis of scRNA-seq vs. those detected in Xenium in situ for TCRs. “Patient-matched” refers to correlations between Xenium samples and scRNA-seq samples originating from the same patient at the same clinical time point where “patient-mismatched” refers to correlations between different patients or different clinical timepoints. B) Correlations of V-gene usage between scRNA-seq vs. Xenium for IG-genes. C) Correlation of T cell “pseudo-clone” sizes with paired B cell “pseudo-clone” sizes as a function of PD-1 inhibitor exposure of the originating tumor tissue and degree of spatial co-clustering between those T and B “pseudo-clones”. D) Representative images of tumor and lymph node samples from patient su001 highlighting a pair of co-clustered “pseudo-clones” and a pair of unclustered “pseudo-clones”. Green lines represent T/B cell clones within 20µm of each other. E) Fraction of TCRα “pseudo-clone” pairs present in the draining lymph node samples that are also present in the respective tumor sample of that patient. All lymph node samples were collected post-PD-1 inhibitor treatment. Patient su005 had no remaining archived pre-PD-1 inhibitor tumor to be analyzed. F) Fraction of TCRβ pairs present in both draining lymph nodes and tumors. G) Normalized gene expression by patient and tissue compartment for T cell clones that are present in both lymph nodes and tumor (shared) vs. those present in a single compartment (exclusive). H) Schematic diagram of a proposed mechanism of antigen cross-presentation by B cells to CD8+ T cells leading to a cycle of tumor clearance and generation of additional tumor neo-antigens.

To examine functionally relevant T- and B-cell interactions, we analyzed spatial transcriptomics data for co-clustering of T- and B-cell clonotype pairs--as defined by the dominant V-gene usage--using neighborhood enrichment analysis, which calculates the degree of spatial co-occurrence between selected cell types (Methods, Palla et al., 2022). We then examined the sizes of T and B clonotypes in our samples as a function of spatial co-localization. As expected, T- and B-cell clones with no spatial co-localization lacked any correlation in clonotype sizes (**Figure 3C**). Interestingly, specifically in aPD-1-treated specimens, we observed significant correlation between B-cell clone size and the number of paired, spatially co-localized T-cell clones. This suggests that PD-1 blockade may promote the expansion of interacting T and B cells.

Previous studies have highlighted the importance of tertiary lymphoid structures (TLSs) in successful responses to PD-1 inhibitors (Meliante et al., 2023, Cristescu et al., 2018, Sautès-Fridman et al., 2016, Helmink et al., 2020, Petitprez et al., 2020, Carbrita et al., 2020). The presence of TLSs could potentially confound our analyses of BCR diversity within the tumor microenvironment. To address this issue, we evaluated hematoxylin and eosin (H&E) staining of our specimens, and while we observed tumor-infiltrating lymphocytes, there were no clearly formed TLSs evident in any of our samples (**Extended Data Figure 4**). This suggests that BCR clonotype diversity represents an important phenomenon independent of histologically distinguishable ectopic lymphoid structures. Given the absence of TLSs within the tumors, we hypothesized that regional draining lymph nodes may also be a critical site for the development of the T/B-cell interactions that we observe in our data.

In two patients, tissue samples from both post-treatment tumors and the tumor-draining lymph nodes were available for analysis. We examined the presence of specific T- and B-cell clonotype pairs in tumors and regional lymph nodes. We find that T/B-cell clonotype pairs that were co-localized in both tumor and lymph nodes were approximately 2-10 times more likely to be in both locations compared to non-colocalized clonotypes, which tended to be found exclusively in either tumor or lymph node (**Figure 3D-F**). This overlap was more pronounced in patient su001 who experienced excellent long-term response and remission of disease compared to patient su005 whose disease was progressing at last available follow-up.

We then examined the phenotypes of these “shared” clonotypes enriched in both tumor and lymph node, which appeared to be trafficking between these sites. In lymph node samples, T cells from these shared clonotypes exhibited higher expression of the proliferation marker *MKI67* compared to other cells (**Figure 3G**). *CD69* expression, an indicator of recent activation, was observed in shared T-cell clonotypes in both patients; however, it was only significantly elevated in the tumor specimen of su001, who achieved remission. Finally, *ENTPD1* (encoding CD39, a marker associated with tumor-reactive T cells) was elevated only in the tumor specimens of su001. These findings highlight the nuanced inter-patient differences in PD-1 inhibitor responses and suggest a model by which B-cell clones support the expansion and activation of specific T-cell clones in lymph nodes leading to more effective tumor clearance (**Figure 3H**). The inter-patient variation in how nodal clonal interactions turn into a successful tumor-reactive T cell response deserves further investigation.

While direct cytotoxic activity against tumor cells is driven by CD8+ T cells, various modes of support for these antigen-specific CD8+ T cells have been demonstrated. Recent studies have identified an immune cell triad of CD4+ T cells, CD8+ T cells and dendritic cells that coalesce to reprogram CD8+ T cells for tumor clearance (Espinosa-Carrasco et al. 2024). Given that B cells, like dendritic cells, function as professional antigen-presenting cells, we hypothesized that the relationship we observe between B cells and CD8+ T cells may also be influenced, in part, by a supportive, co-localized CD4+ T cell interaction. Our co-embedded spatial transcriptomics dataset, incorporating aPD1-exposed tissue samples, uniquely enables the study of this phenomenon.

To quantitatively assess the presence of cell cluster triads, we defined a metric we call ‘*Triad Occurrence*’ (**Extended Data Figure 7A**, Methods) that measures the co-occurrence probability of having a third cell type in proximity to an existing proximal cluster pair. As a proof-of-concept, we first tested this measurement on our malignant cell clusters since these clusters of tumor cells exist in well-structured tumor islands and may represent phylogenetically related daughter cells. Compared to a triad of three cell types with no expected co-localization, these malignant cell clusters showed a significant propensity to co-localize, validating the utility of this analysis to detect the occurrence of cell cluster triads (**Extended Data Figure 7B-C**).

Next, we tested CD4+ T cell/Activated CD8+ T cell/Dendritic cell triads, which showed increased intratumoral co-clustering in agreement with recent studies (**Extended Data Figure 7D**, Espinosa-Carrasco et al. 2024). In our prior study, we noted parallel development of follicular helper T cells (Tfh) and CD8+ T cells. Given our observations of B cell involvement in T cell activation in regional lymph nodes, we asked whether there might be a B cell/Tfh/Activated CD8+ T cell triad present in our patient samples. Indeed, we observe co-occurrence of this B cell/Tfh/Activated CD8+ T cell triad, but interestingly, specifically in regional lymph node samples, suggesting that a substantial portion of this interaction happens in the tumor-draining nodes (**Extended Data Figure 7E-F**).

### Trajectory of Induced BCR Diversity Correlates with Post-Immunotherapy Survival in Multiple Cancer Types

To determine whether the correlation between BCR diversity and survival is BCC-specific or represents a more generalizable phenomenon following checkpoint blockade, we aggregated data from published studies that include pre- and post-immunotherapy tumor sequencing and clinical outcomes. Identified studies meeting our criteria included head and neck squamous cell carcinoma, melanoma, and glioblastoma which we combined with our existing BCC dataset (Yost et al. 2019, Riaz et al. 2017, Cloughesy et al. 2019, Liu et al 2021). We extracted BCR clonotypes from these studies and categorized the patients based on the induction of BCR diversity following checkpoint blockade using the same criteria established in our earlier analysis. Across this aggregated dataset which included cancers of various embryonic origins, induced BCR diversity was associated with better clinical response (**Figure 4A**). In the cohort of glioblastoma patients where progression-free survival data was available, higher BCR diversity was associated with prolonged progression-free survival specifically in patients treated with PD-1 inhibitor in the neoadjuvant setting whereas their treatment-naïve counterparts with a scheduled, adjuvant course showed no significant difference (**Extended Data Figure 8**). In the combined cohort of 93 patients across these four cancer types, overall survival was significantly better in patients exhibiting induced BCR clonal diversity following checkpoint blockade (**Figure 4B**) (HR = 0.46 95% CI 0.245882 −0.8678724, log-rank p = 0.0093).

**Figure 4:**
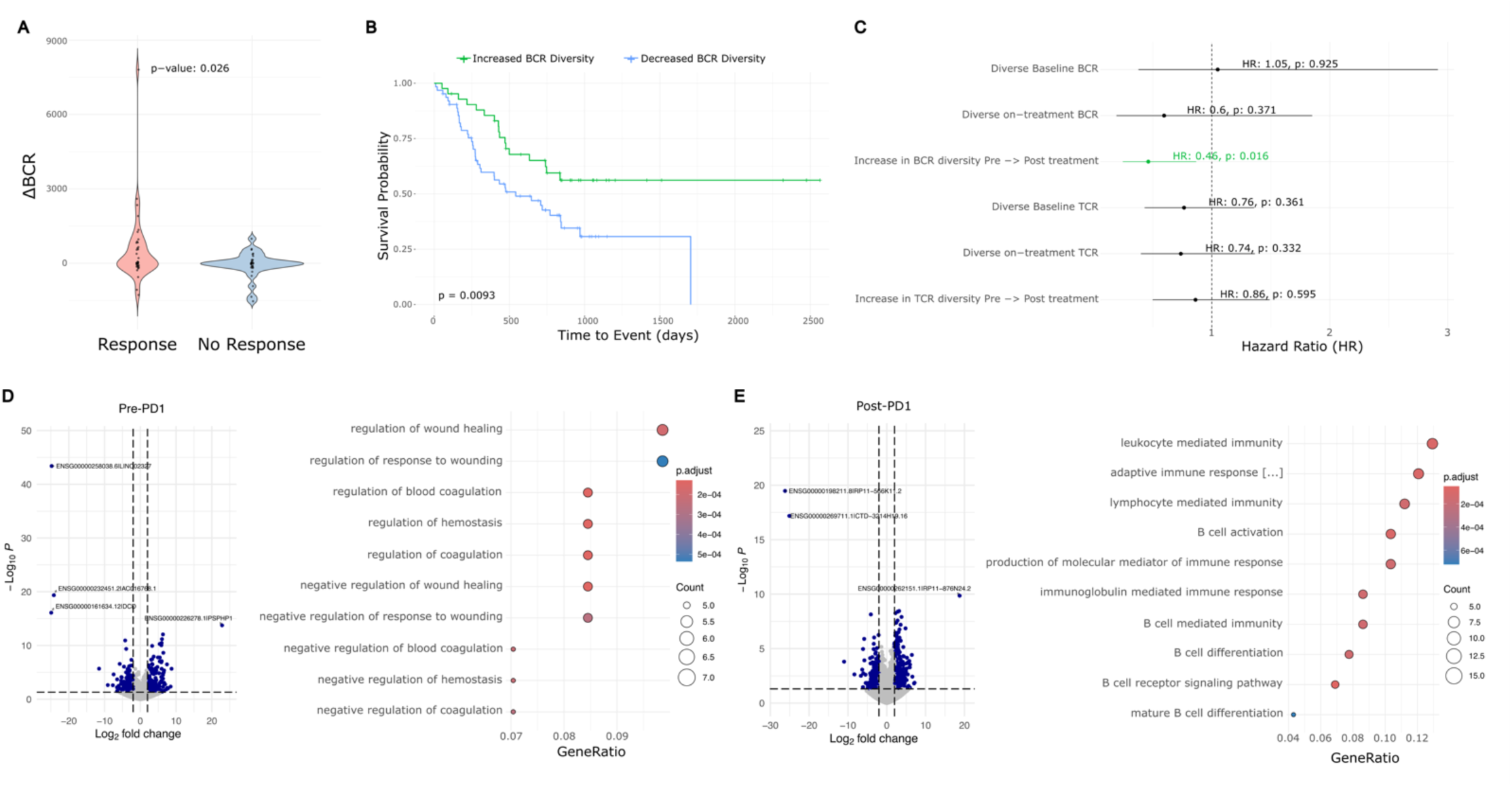
Meta-analysis of BCC, glioblastoma, melanoma, and HNSCC reveals a general prognostic role for aPD1-induced BCR diversity. A) ΔBCR clonotype count from pre-PD1 inhibitor to post-PD1-inhibitor of patients with progressed tumors vs. non-progressed tumors. B) Kaplan-Meier curve of overall survival for aggregate cancer patients stratified by induced BCR expansion from PD-1 inhibitor. C) Hazard ratios of overall survival for aggregate cancer patients by BCR or TCR clonotype. An increased clonotype diversity is defined by a greater number of clonotypes detected in post-PD1 treated tumors compared to prior. Diversity in the case of static measurements at baseline or on-treatment are defined as an above-median BCR or TCR clonotype count. D) Left: Volcano plot of differentially expressed genes in pre-treatment tumors that undergo induced BCR expansion vs. those without. Right: GO-terms enriched in pre-treatment tumors that undergo induced BCR expansion vs. those without. E) Left: Volcano plot of differentially expressed genes in post-treatment tumors that undergo induced BCR expansion vs. those without. Right: GO-terms enriched in post-treatment tumors that undergo induced BCR expansion vs. those without

Prior studies have suggested the potential prognostic importance of static BCR clonotype measurements (Valpione et al 2022, Singh et al 2022, Petitprez et al 2020), however, others have failed to detect significant associations (Damsky et al. 2019). We therefore sought to evaluate this in our aggregated dataset. Consistent with previous reports, we observed a trend suggesting protective effects of higher static BCR diversity, particularly when measured in treatment-exposed tumors (**Figure 4C**); however, in our multi-cancer cohort, the dynamic trajectory of induced BCR clonotypes following checkpoint blockade was more directly related to survival outcomes and was the only criterion that reached statistical significance.

### A Machine-Learning Model Predicts the Trajectory of BCR Clonotype Diversity from Pre-Treatment Tumor Gene Expression

Although induced BCR diversity is a potentially important prognostic factor for immunotherapy response, measuring it in routine practice is challenging. Calculating induced BCR diversity requires adequate tumor specimens for sequencing from both the pre-treatment tumor as well as from post-treatment tumors. While obtaining multiple sequential samples is practical in certain circumstances--such as when tumors are easily accessible (*e.g.*, cutaneous neoplasms), or when neoadjuvant checkpoint blockade precedes a planned surgical resection attempt--in most cases sequential sampling would expose the patient to additional, unplanned procedural risks. Therefore, it would be beneficial to understand whether this information could be derived solely from the baseline data from pre-treatment tumors, thus providing more timely data for clinical decision making while simultaneously avoiding the risks of additional tissue sampling.

To address this, we re-annotated our multi-cancer cohort based on the patients’ eventual change in BCR diversity. Interestingly, pre-treatment tumors that respond to PD-1 therapy with an induced increase in BCR diversity were enriched for genes related to the coagulation cascade (**Figure 4D**). While studies have linked coagulation with the function of PD-1 inhibitors, to our knowledge this is not a known positive prognostic signature (Sato et al 2019). High endothelial venules – specialized post-capillary structures known for their presence in secondary lymphoid organs – are known to enhance lymphocyte recruitment (Milutinovic et al. 2021). These data pose a possibility that there may be specific vascular structures or signatures that prime the tumor for improved lymphocyte turnover, but this requires further, separate investigation. As expected, the post-treatment tumors with increased BCR diversity were enriched for gene expression relating to B cell function including “B cell activation”, “B cell receptor signaling pathway”, and “Lymphocyte/leukocyte mediated immunity” (**Figure 4E**).

We hypothesized that these differentially expressed genes may be diagnostically important in identifying which tumors are primed for induced BCR diversity. To leverage this possibility, we developed a machine-learning model using this gene expression profile as input to categorically classify potential BCR responses to checkpoint blockade. We trained this neural network model on the 117 identified differentially expressed genes, achieving a 92.3% accuracy on a 20% withheld set of pre-treatment tumor specimens (Methods, **Supplemental Data Table 3**). This result suggests that baseline tumor gene expression may contain sufficient information to predict induced BCR expansion.

There are however limitations to this approach that need to be addressed. First, since the data in the training and validation sets originated from the same cohort, the validation does not account for real-world variability introduced by varying sample collection protocols and standards. Second, while induced BCR diversity is mechanistically interesting, it is not clinically actionable absent a direct link to treatment outcomes. To address both limitations, we subjected the model to a more challenging two-step inference problem: predicting clinical outcomes using only baseline tumor RNA sequencing data from a completely independent dataset. The model was completely naïve to this new dataset, simulating the challenges of applying the model to a new cohort. Additionally, to ensure that the model was not simply identifying a non-specific positive prognostic signature but rather one specific to checkpoint blockade, we included a control cohort of patients treated with non-immunotherapy approaches including chemotherapy, radiation, and targeted molecular therapies who had no opportunity to induce a checkpoint inhibitor-specific mechanism of response. We identified cohorts of melanoma patients in published data and The Cancer Genome Atlas Program meeting these criteria and proceeded with modeling on these cohorts (Liu et al. 2019, Gide et al. 2019, Weinstein et al. 2013).

Among the melanoma patients undergoing checkpoint blockade, those whose baseline tumors were predicted by the model to increase BCR diversity following treatment showed higher proportions of clinical responses (including marginal, partial, and complete responses), while those predicted not to have an increase in BCR diversity had higher proportions of progressive disease (**Figure 5A**). Additionally, patients with predicted increases in BCR diversity had both superior progression-free survival (log-rank *p*=0.00038) and overall survival (log-rank *p*=0.00046) compared to their counterparts in whom the model predicted no increase in BCR diversity (**Figure 5B-C**). Supporting the checkpoint inhibitor-specific function of the model, patients who underwent treatment with non-PD-1 modalities did not show stratification by survival outcomes based on the model’s predictions (log-rank *p*=0.53 and *p* = 0.60 for PFS and OS, respectively).

**Figure 5:**
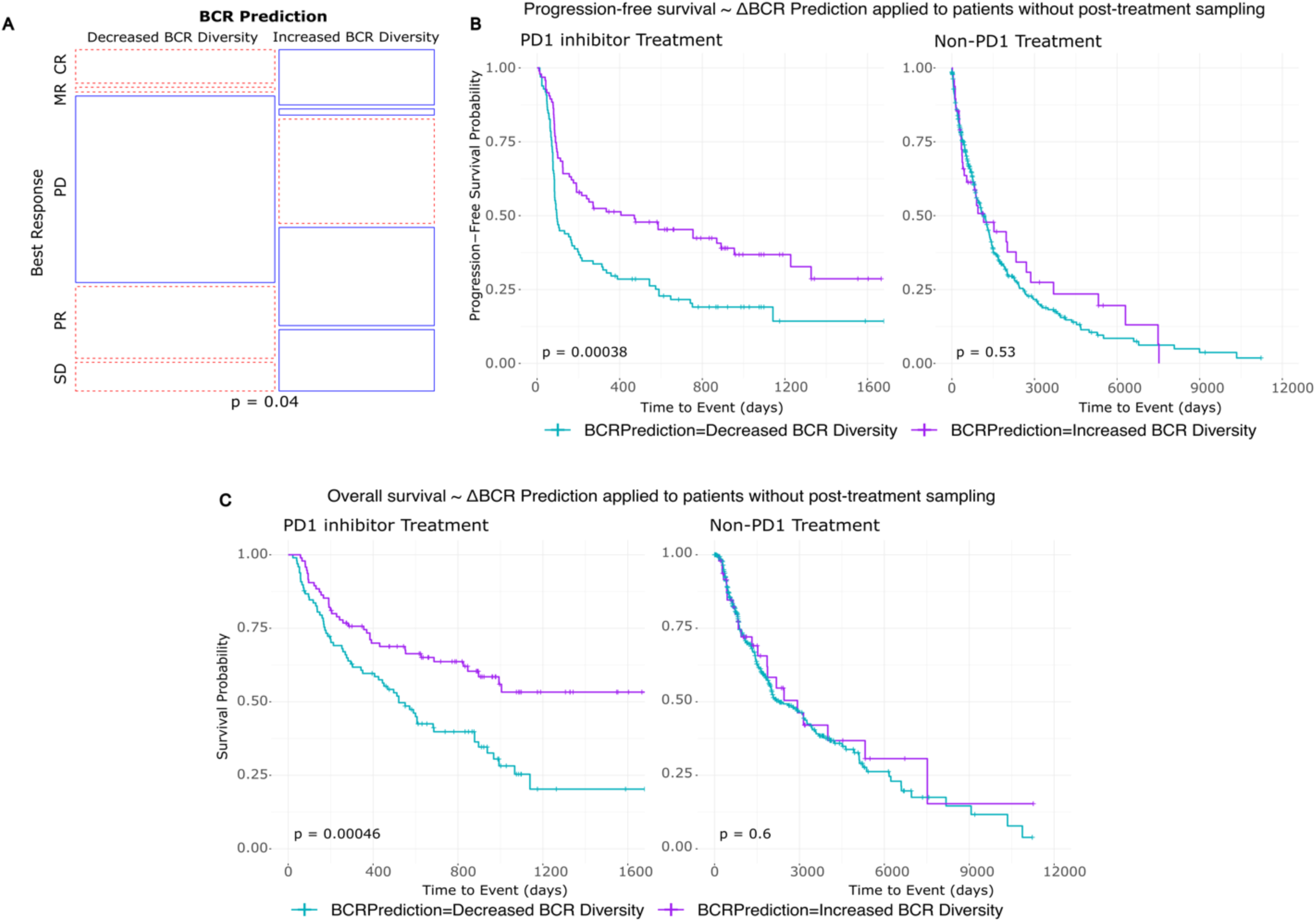
Pre-treatment tumor RNA sequencing can be leveraged to model aPD1-induced BCR diversity and clinical outcomes. A) Prediction model applied to a new melanoma patient cohort. Mosaic plot of clinical response by RECIST stratified by model prediction of induced BCR clonotype expansion. BR: Best response, CR: Complete response, MR: Marginal response, PD: Progressed disease, PR: Partial response, SD: Stable disease. B) Kaplan-Meier curve of progression-free survival stratified by model prediction of induced BCR clonotype expansion in melanoma patients treated with PD1 inhibitor (left) and other, non-PD1 therapies (right). C) Kaplan-Meier curve of overall survival stratified by model prediction of induced BCR clonotype expansion in melanoma patients treated with PD1 inhibitor (left) and other, non-PD1 therapies (right).

## DISCUSSION

Immunotherapy has emerged as a leading treatment option for many advanced cancers. Its potential is evident in the positive clinical responses and cases of long-term remission that can be induced by checkpoint blockade therapies. The promise, however, is juxtaposed against many cases of inadequate response and an incomplete understanding of the mechanisms underlying treatment failure. The effectiveness of immunotherapy on a patient-by-patient basis arises from an ecosystem of interactions between the tumor, the patient’s immune state, their environment, among other factors (Chen et al. 2017).

In this study, we collected data under real-world circumstances to pair clinical response with underlying biology. We find that one potential outcome of checkpoint blockade therapy is the induction of BCR clonotype diversification. This diversification appears to support the activation and proliferation of CD8+ T cells in regional lymph nodes that then traffic to the tumor to enhance tumor clearance. The successful induction of BCR clonotype expansion is closely linked to patient outcomes, such as overall survival, not only in our cohort of BCC patients but additionally in cohorts of multiple cancer types. These findings support previously hypothesized mechanisms of B-cell anti-tumor activity through their unique abilities to both clonally expand and present antigen (Nielsen and Nelson, 2012) and agree with other studies highlighting the importance of tumor-infiltrating CD8+ T cells in improved cancer survival (Guo et al. 2023). Agents that promote B cell diversification, such as the CD40 or OX40 pathways, may directly impact this mechanism.

Despite these encouraging findings, our study has limitations including its retrospective nature and focus on specific malignancies. Future studies on prognostic use should be validated in a prospective manner, and studies with larger populations and more diverse tumor types would help clarify the generalizability of these mechanistic findings. Nevertheless, we hope this study advances the field in seeking better prognostic tools for cancer patients undergoing checkpoint blockade and guides improvements in immunotherapies for non-responders to current regimens (Nielsen and Nelson, 2012, Tonkunaga et al. 2019).

## Materials and Methods

### Human subjects

As previously described (Yost 2019), data collected in this study was approved by the Stanford University Administrative Panel on Human Subjects in Medical Research (IRB protocol number 18325). Our institution complies requirements for protection of human subjects including 45 CFR 46. All participants provided written informed consent. The patients included in this follow-up had histologically-confirmed advanced or metastatic basal cell carcinoma not suitable for surgical resection at the time of enrollment. Patients underwent treatment with pembrolizumab (200mg every 3 weeks) or cemiplimab (350mg every 2 weeks). Exclusion criteria included prior exposure to checkpoint blockade agents, systemic immunosuppression, exposure to radiotherapy, and use of any other anti-cancer agents within 4 weeks of first biopsy for scRNA-seq. Patient su003 had a distant >50 year prior history of radiation for acne at an unrelated site, patient su004 had adjuvant radiation for a squamous cell carcinoma at a unrelated site two years prior to enrollment, and patient su006 received unspecified radiation for a childhood medulloblastoma > 50 years prior to enrollment. The remaining patient did not undergo radiotherapy prior to enrollment. The medical records for these patients were accessed in November 2023 to supplement long-term survival data not available at the time of prior data publication. Where there was a clearly documented date of death, this was used as the endpoint for survival analyses. In the absence of a date of death, the last confirmed medical contact in our medical record was used as the date of censorship. Two separate investigators were responsible for updating this clinical record with the second investigator blinded to any molecular data analyses at the time of medical record review to reduce bias.

### Tumor collection, library preparation, and sequencing

The in-depth protocol for the collection and processing of human patient samples is as previously described (Yost et al. 2019). In brief, tissue samples were collected during clinical visits at Stanford Hospital and Clinics. Written consent was obtained prior to each tissue collection. The local area of tissue sampled was marked, photographed, and anesthetized with lidocaine. The subsequent tissue processing was performed as previously described for each downstream application. Biopsy specimens were obtained using a 4mm punch biopsy device. Wound care was performed as clinically indicated for the collection sites.

For single-cell RNA sequencing, the samples were processed using the 10x Single Cell Immune Profiling Solution Kit according to the manufacturer’s instructions. The resulting libraries were sequenced on either an Illumina NextSeq or HiSeq 4000 to a minimum sequencing depth of 25,000 reads per cell with the following read lengths: Read1 – 26, i7 index: 8, Read 2: 98. The sequencing reads were aligned to the GRCh38 reference genome and quantified using Cellranger count (10x Genomics, Version 2.1.0). For clinical care specimens, biopsy sites were prepared and anesthetized in the same manner. Specimens were collected through a tangential biopsy technique with a curved sterile blade (Derma Blade Shave Biopsy Instrument), punch biopsy tool, or scalpel depending on the context of specimen collection. Tissues were immediately submerged in formalin 10% and then subsequently embedded in paraffin. Tissue sections were cut as required for clinical diagnosis and the remaining blocks were preserved in long-term storage. These tissue blocks were requested at the time of this study for additional analysis as described below.

### Single cell RNA sequencing analysis

ScRNA-seq data from Yost et al., 2019 were obtained from the Gene Expression Omnibus and the supplementary data of the publication. These data were re-analyzed using Seurat (Version 5.0.3). Identical to prior cutoffs, cells with less than 200 genes detected or greater than 10% mitochondrial RNA content were excluded. Cell cluster annotations were performed based on the expression of known marker genes as previously reported, summarized below.

| Cell Type | Marker Genes |
| --- | --- |
| T cells | CD3G, CD3D, CD3E, CD2 |
| CD8<sup>+</sup> T cells | CD8A, GZMA |
| CD4<sup>+</sup> T cells and T<sub>reg</sub> cells | CD4, FOXP3 |
| NK cells | KLRC1, KLRC3 |
| B cells | CD19, CD79A |
| Plasma cells | SLAMF7, IGKC |
| Macrophages | FCGR2A, CSF1R |
| Dendritic cells | FLT3 |
| Plasmacytoid dendritic cells | CLEC4C |
| Fibroblasts | COL1A2 |
| Myofibroblasts | MCAM, MYLK |
| Cancer-associated fibroblasts | FAP, PDPN |
| Malignant cells | EPCAM, TP63 |
| Endothelial cells | PECAM1, VWF |
| Melanocytes | PMEL, MLANA |

ScRNA-seq data was analyzed as previously described (Yost et al. 2019). Cell clustering was performed with Seurat using the same cell lineage markers previously described. The cluster identities were largely in agreement to prior, with the updated analysis showing differences only in subclustering of malignant cell clusters and the identification of a small plasma blast population. Receptor-ligand and signaling pathway relationships between single cells were analyzed using Cellchat (Version 2.1.0, Jin et al., 2021). Diffusion maps and pseudotime analyses were created with the R package Destiny (Haghverdi et al. 2015, Version 3.16.0).

### TCR and BCR clonotypes

T-cell receptor and B-cell receptor repertoires were identified using TRUST4 (Song et al. 2021, Version 1.0.13). TRUST4 was run on the scRNA-seq FASTQ files with default parameters to extract V(D)J sequences from unassembled reads. These clonotypes were defined based on V/J gene usage and CDR3 sequences. A unique clonotype was considered only if there were three or more supporting reads. Analysis of this dataset required the use of Stanford’s “Sherlock” high performance computing cluster. Further downstream analyses were performed using the R programming language (R Development Core Team, http://www.r-project.org/) and using the R package Immunarch (https://immunarch.com/, Version 0.9.1).

### Correlation analysis of BCR diversity-associated gene expression

To identify genes whose expression levels are associated with BCR clonotype counts, we performed a correlation analysis using the scRNA-seq dataset. To additionally understand how individual cell clusters were contributing to this gene expression correlation and whether there was a dominant cell cluster which might be influenced by tissue BCR diversity, we implemented a cluster elimination strategy as a form of regression analysis, where cell clusters were systematically excluded and the correlations recalculated to assess each cluster’s impact on gene-BCR correlations.

Gene expression counts were obtained from the log-normalized counts in the Seurat data. BCR clonotype counts derived from TRUST4 analysis of the tissue specimens were annotated in the metadata of the Seurat object. We computed Pearson correlation coefficients between the expression of each gene and the BCR clonotype counts present in the tissue specimen.

The cluster elimination strategy was implemented by creating a new Seurat object that systematically excluded each cell cluster from the dataset and re-performed the correlation calculations described above. We defined the change/delta in correlation as the difference between the Pearson correlation of the data with the specific cell cluster of interest excluded vs. that of all the cell clusters without exclusion. Visualization of this data was performed using ComplexHeatmap (Version 2.18.0).

### Cell cluster triad analysis

Cell clusters in the spatial transcriptomics dataset were annotated using the nearest-neighbor approach in the ENVI latent space described separately. The CD4+ T cell population was subdivided into a CXCR5+ population based on detectable probe counts that we refer to as Tfh cells. Specifically for this analysis we propose a new metric that quantified the frequency of a third co-localizing cell type in proximity to a co-localizing cluster pair of interest. We term this metric the Triad Occurrence as defined below. The distance threshold d is set to be the approximate diameter of an average cell. The p-value was calculated by generating an empiric null distribution over 10,000 iterations and comparing the true observed value to this null distribution.

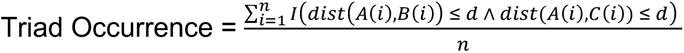

*n*: *Total occurences of cluster A* (*origin cell*)
*d*: *distance threshold*. *In this case* 10µm
*dist*<*A*(*i*), *B*(*i*)): *distance between the i*
− *th occurence of cell belonging to cluster A and the nearest cell belonging to cluster B*
*dist*<*A*(*i*), *C*(*i*)): *distance between the i*
− *th occurence of cell belonging to cluster A and the nearest cell belonging to cluster C*
*I*(*x*): *Indicator function*. *I*(*x*) = 1 *if x is true and I*(*x*) = 0 *if x is false*

### Statistical analysis

Time to event analyses (PFS, OS, BCC/SCC risk) were performed using the Kaplan-Meier method. Hazard ratios were calculated using the Cox proportional hazards model. Contingency tables were analyzed using the chi-square test. P-values are as reported in the main text with the default reporting being the p-value from the log-rank test unless otherwise stated.

### Realignment of RNA sequencing differential gene expression analysis

FASTQ sequences obtained from GEO were realigned to the GRCh38 reference genome using HISAT2 (Version 2.1.0, Kim et al. 2019) using default parameters. Gene quantification was performed using Stringtie (Version 2.2.1, Pertea et al. 2015). Differential gene expression was conducted using DEseq (Version 1.12.3, Love et al. 2014). Genes with an adjusted p-value < 0.05 and an absolute log2 fold change > 1 were considered significantly differentially expressed. Where FASTQ files were not available, the aligned counts from the referenced datasets were used directly.

### Machine Learning of baseline tumor characteristics for BCR predictions

To Predict BCR clonotype expansion based on baseline tumor gene expression, we developed a neural network model using the Keras (Version 2.13.0) library in R. Linear regression-based models were unable to generate predictions that exceeded random chance. The input features were the 117 differentially expressed genes identified from pre-treatment tumors.

The dataset was split into 80% training and 20% validation sets using stratified random sampling to maintain proportional representation. The neural network architecture was designed to minimize complexity and therefore avoid overfitting. The final architecture consisted of:

- An input layer with 117 units corresponding to the input features.
- A hidden layer with 64 units and ReLU activation.
- A dropout layer with a rate of 0.5.
- A second hidden layer with 32 units and ReLU activation.
- A dropout layer with a rate of 0.5.
- An output layer with 1 unit and linear activation for regression output.

The model was compiled using the Adam optimizer and used a mean squared error loss function. Training was performed over 800 epochs and early stopping implemented based on validation loss. Model performance was evaluated using the root mean squared error on the categorical assignments of BCR clonotype change and the coefficient of determination on the validation dataset.

### Xenium In Situ panel design

Genes were selected for Xenium In-Situ based on multiple criteria. First, we automatically included all genes from our prior study that were required for cell type or cell state identification. Next, we sought the assistance of the 10x design team to assist with the design of custom probes specific to individual V(D)J genes. These probes were validated internally by 10x Genomics. We next identified the most differentially expressed genes present in our prior scRNA-seq dataset and included the maximum number of these genes in order of degree of differential expression up to the maximum allowed capacity of 480 genes at the time of design. The final design was assessed using the custom design panel from 10x genomics. Two panels – “Human Skin (aging)” and Human Skin (normal/inflamed)” were used as design references in the quality control. 475 out of the 480 selected genes were within the recommended expected expression range to avoid optical crowding, however, given our assessment of the expression in scRNA-seq data, all 480 were retained for the final panel design.

### Xenium In Situ

Xenium In Situ Gene Expression assays were run according to the manufacture’s (10x Genomics) instructions. Custom probe kits and accompanying reagents were ordered from 10x Genomics and stored at their recommended temperature until use, in our case within 6 months.

A 3 mm diameter core was punched from the region of interest (ROI) of each FFPE donor block and placed into its designated position within the recipient block. Cores were arranged in a 2×4 or 2×5 sample array to align with the available Xenium slide area. The recipient block, containing the specimens, was placed into a paraffin embedding station to fill any gaps and secure the cores. Five-micron sections of the TMA block were then cut using a microtome and mounted onto slides. Quality control (QC) was performed by staining and examining test sections to confirm that the cores were intact and properly oriented. Care was taken to ensure that the microarray grid did not interfere with the slide fiducials during Xenium slide preparation. Microarrays were sectioned directly onto the Xenium slide in five-micron sections immediately prior to slide processing. Following sectioning, the Xenium slides were air-dried under a fan for 30 minutes and then baked at 42°C for 3 hours. The slides were subsequently stored overnight in a desiccated container at room temperature.

Deparaffinization of the tissue slides was performed as follows: 1) Incubation at 60C for 2h. 2) Cooling tissue to RT for ∼7min. 3) Immersion in xylene × 10 min x 2. 4) Immersion in 100% ethanol x 3 min x2. 5) Immersion in 96% ethanol for 3 min x2. 6) Immersion in 70% ethanol x 3 min. 7) Immersion in nuclease-free water for 20s. 8) Wash with 1x PBS.

Decrosslinking of the tissue slides was performed as follows: 1) Incubation of Decrosslinking buffer (see manufacturer’s instructions) at 80C x 30min then 22C for 10 min. 2) Wash with 1x PBS-T x 1 min x 3.

Custom probes were resuspended in TE buffer according to manufacturer instructions immediately prior to use. Probe hybridization was performed overnight at 50C. Post-hybridization washes were performed with PBS-T at RT then again at 37C x 30miin. The probe ligation reaction was performed at 37C for 2h followed by PBS-T washes at RT. Amplification was performed at 30C for 2h and followed by TE buffer washes at RT. Autoflorescence quenching was performed to chemically mitigate background florescence according to the manufacturer instructions. Prepared slides were loaded onto the Xenium Analyzer instrument running Xenium Onboard Analysis v2.0.

To improve the chances of capturing accurate cell boundaries, all Xenium slides were additionally run with multi-modal cell segmentation. This kit contains a variety of cell stains including DAPI for nuclear staining/segmentation, ATP1A1, E-Cadherin, and CD45 for establishing membrane boundaries, 18S ribosomal RNA labels to detect the cytoplasm, and alphaSMA/Vimentin for interior protein staining. The default isotropic nuclear expansion distance is set as 5 μm. Standard H&E staining was performed immediately after the decoding and imaging. Briefly, the post-run slide was immersed in Quencher Removal Solution (10x Genomics) and then washed with Mili-Q water. Staining was performed with Hematoxylin, Bluing solution, and then Eosin according to manufacturer instructions.

Critical data files of the output were exported off the Xenium Analyzer Instrument. Image registration and initial data exploration was performed in Xenium Explorer before being exported to other community-based, open-source methods as separately described.

Kits and reagents from 10x Genomics were used for the preparation of Xenium slides and the Xenium Analyzer run, including: Xenium Slides & Sample Prep Reagents (PN-1000460), Xenium Decoding Consumables (PN-1000487), Xenium Decoding Reagents (PN-1000461), and Xenium Cell Segmentation Staining Reagents (PN-1000661).

### Neighborhood enrichment analysis

Neighborhood enrichment was run for the Xenium data using Squidpy (Version 1.6.0, Palla et al. 2022). Cells expressing V-genes were categorized by the highest-expressing V-gene within their segmented cell boundary. These assignments were then used as surrogate cell clusters using the sq.gr.spatial_neighbors function within Squidpy. The neighborhood enrichment analysis calculates an enrichment z-score based on the proximity on the connectivity graph of these clonotype assignments. A z-score of 2 was used as a cutoff for spatially significant enrichment for downstream analyses.

### BCC cohort survival analysis

Patients su001 to su008 from Yost et al., 2019 were included in the analysis. Patient su009 to su013 were excluded due to the absence of BCR data (T-cell isolates only were obtained). Patient su004 who lacked detectable immunoglobulin clones in both pre- and post-PD-1 treatment samples was classified as a BCR non-expanded patient. To minimize noise contributed by non-specific BCR calls and to focus the analysis on clones more likely to be functionally relevant in the tumor, a minimum threshold of 3 counts per clones was established to define a significant BCR clone. This represented the same threshold cutoff used in our prior study.

Patient were categorized into two groups based on the dynamic change from pre- to post-PD-1 treated tumors.

BCR Expanded: Patients exhibiting an increase in BCR clone counts in post-PD-1 treatment tumors compared to pre-PD-1 treatment tumors.

BCR non-Expanded: Patient exhibiting no change or decrease in BCR clone counts in post-PD-1 treatment tumors compared to pre-PD-1 treatment tumors.

The change in BCR clone counts was defined as follows:

ΔBCR=BCRpost-treatment−BCRpre-treatment

Survival times were computed from the date of first PD-1 inhibitor exposure until either the date of death – when available – or the last follow-up date for censored observations. Kaplan-Meier curves were generated using a combination of the R packages survival (Version 3.5.8), Survminer (Version 0.4.9), and plotted with either ggplot2 (Version 3.5.0) or using Graphpad Prism 10. Survival distributions were compared between groups using the log-rank test. Hazard ratios and 95% confidence intervals were estimated using the Cox proportional hazards regression model to assess the impact of BCR clonal expansion on overall survival.

### ENVI cointegration of scRNA-seq and Xenium

ENVI (Version 0.3.6, Haviv et al. 2024) was run using the CPU-based computation paramenters given the large memory requirements of our data-set. Dependencies for this analysis were installed according to the instructions recommended in the package documentation. Prior to analysis both scRNA-seq and Xenium data were re-normalized to a target sum of 10,000 counts per cell and log-transformed. To ensure a consistent latent space, ENVI was run with all spatial samples represented in each model initialization. For each subset, the model was trained using the default parameters. Due to memory constraints, the data was batched into 5 random subsets and processed individually with consistent results obtained from each subset.

Dimensionality reduction was performed using Uniform Manifold Approximation and Projection (UMAP) with parameters set to 200 nearest neighbors and a minimum distance of 0.05. Cell type predictions for the spatial transcriptomics dataset were created by identifying the nearest-neighbors with respect to Euclidean distance in the high-dimensional latent space generated by ENVI of known cell cluster assignments in cells originating from the scRNA-seq dataset (Hao et al. 2021, Janesick et al. 2023).

### SpatialData analysis and Visualization

Xenium data were managed using a combination of Xenium Explorer and the python package SpatialData (Version 0.2.0). Xenium Explorer was used for registration of the RNA expression coordinates with the concurrent H&E staining of the same tissue. Given the arrayed format of the tissues, once the coordinates of the individual samples were determined, the data/images for each tissue sample were cropped to the correct areas and stored in Zarr format.

## Data availability

RNA sequencing analyzed in this study was obtained from the Gene Expression Omnibus (GEO), European Nucleotide Archive (ENA), and dbGAP at the following accessions: GSE123813 (Yost et al. 2019), GSE121810 (Cloughesy et al. 2019), phs000452.v3.p1 (Liu et al. 2019, only gene counts accessed), PRJEB23709 (Gide et al. 2019), GSE91061 (Riaz et al. 2018), and GSE179730 (Liu et al. 2021). For the melanoma control dataset, patient sequencing and clinical data were downloaded from the cancer genome atlas Skin Cutaneous Melanoma PanCancer Atlas using cBioportal. Patients with PD-1 inhibitors in their treatments were excluded for the non-PD-1 melanoma control cohort. Xenium data will be deposited and publicly available at the time of publication.

## Acknowledgements

We thank the patients and their families for their generous donations of time and tissue specimens to this study; members of the Chang and Satpathy labs and the department of Dermatology at Stanford for discussions; the department of Pathology at Stanford for assistance with retrieving archived patient specimens; BioChain Institue Inc. for assistance in performing Xenium In Situ experiments. Supported by Parker Institute for Cancer Immunotherapy (A.T.S., H.Y.C.). Y.C. and J.L are supported by the NIH T-32 Institutional Training Grant (2T32AR007422). H.Y.C. is an Investigator of the Howard Hughes Medical Institute.

## Contributions

Y.C., A.L.S.C, and H.Y.C conceived the project. A.L.S.C collected scRNA-seq tumor specimens and provided direct clinical care. K.E.R provided histopathologic analysis of patient specimen and directed procurement of archived patient samples. Y.C., F.A.T, and A.L.S.C performed retrospective review of patient medical records. Y.C. performed spatial transcriptomics experiments and data analysis. Y.C., J.L., A.L.S.C, and H.Y.C guided experimental design and data analysis. Y.C., A.L.S.C, and H.Y.C wrote the manuscript with input from all authors.

---

**Extended Data Figure 1:**
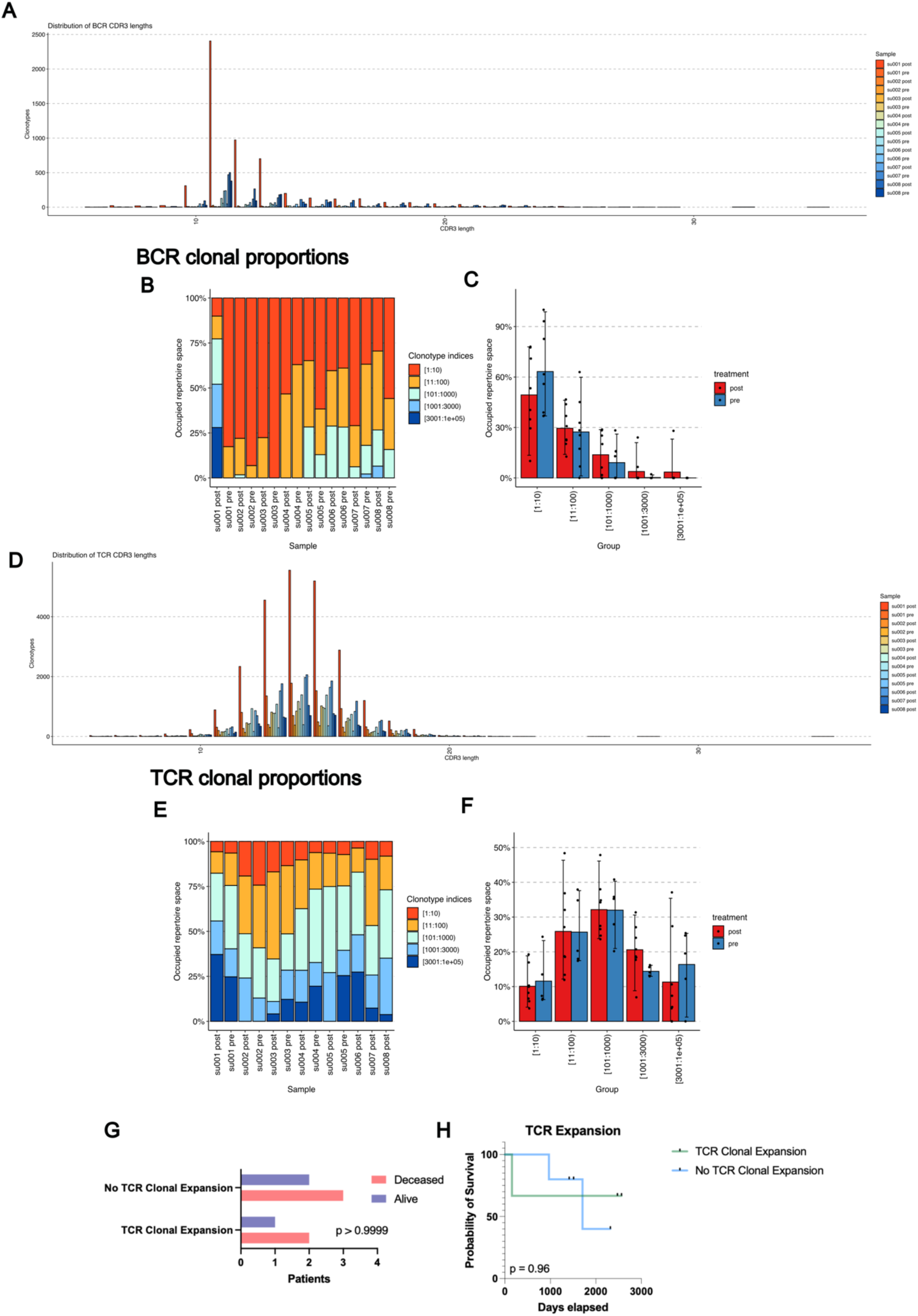
BCR and TCR Clonotype Characteristics. A) Distribution of BCR CDR3 lengths by sample. B) Clonal proportions of BCR clonotypes present in BCC patient specimens expressed as a percent of total. C) Occupied repertoire space of BCR clonotypes present in BCC patient specimens stratified by sample pre vs. post PD1 inhibitor. D) Distribution of TCR CDR3 lengths by sample E) Clonal proportions of TCR clonotypes present in BCC patient specimens expressed as a percent of total. F) Occupied repertoire space of TCR clonotypes present in BCC patient specimens stratified by sample pre vs. post PD1 inhibitor. G) Contingency table of patient status at last clinical follow up vs. the presence of significant clonal expansion in any T cell subset previously described in Yost et al. 2019. H) Kaplan-Meier curve of overall survival for our BCC patient cohort stratified by presence or absence of T cell subset expansion.

**Extended Data Figure 2:**
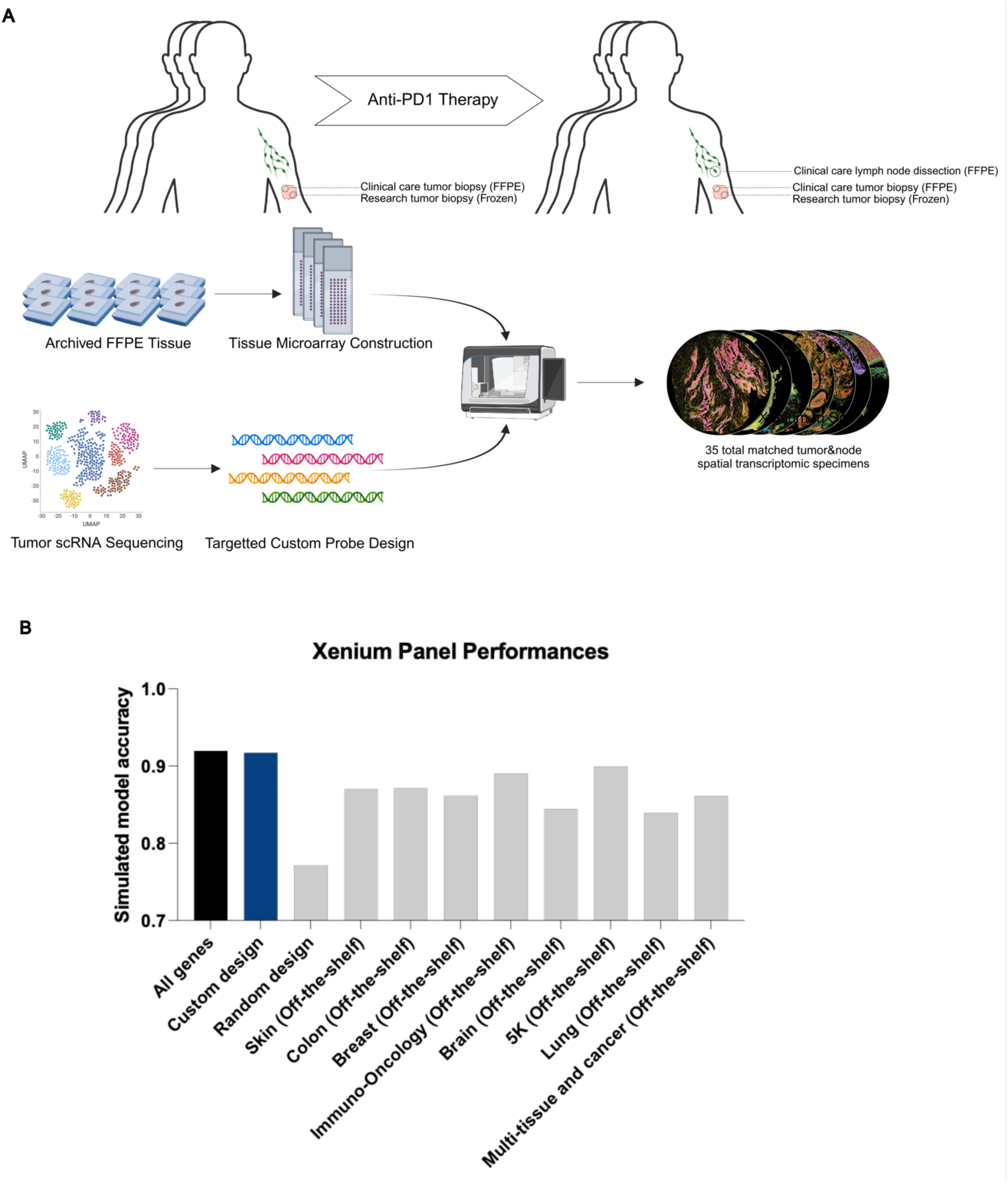
Xenium In-Situ Experimental Design and Probe Design Validation. A) Schematic diagram of Xenium in situ experimental design using archived specimens. B) Model performance of a neural net classifier trained on cell type prediction from scRNA-seq data. A theoretical maximum of model performance was established by allowing full gene expression data to be used in training (black) while all other models were restricted to the genes available in the annotated Xenium panels The custom design (blue) is the 480 gene manually curated gene panel used for our analysis, the random design is a panel of 480 randomly selected genes while all other panels are pre-designed panels offered by 10x genomics.

**Extended Data Figure 3:**
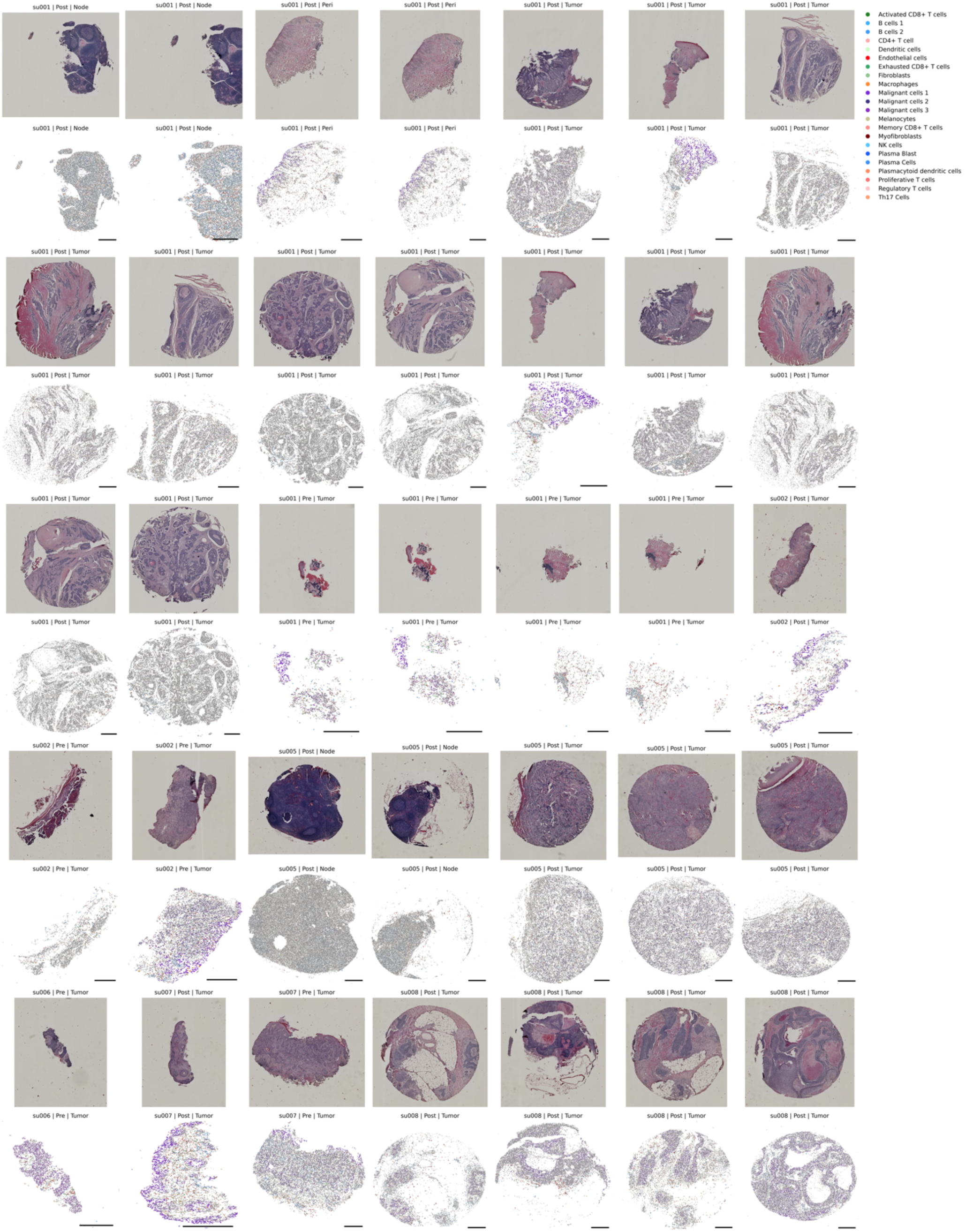
H&E and Xenium In-situ of Primary Patient Specimens. Patient tissue specimens sent for Xenium in situ analysis. Top panels represent H&E staining of the identical tissue section run for Xenium. Bottom panels are spatial cell boundaries (grey) colored by cell type predictions generated through the previously described nearest neighbor approach in the ENVI latent space. Scale bars are 0.5mm. Xenium cells are downsampled to 40% for visualization.

**Extended Data Figure 4:**
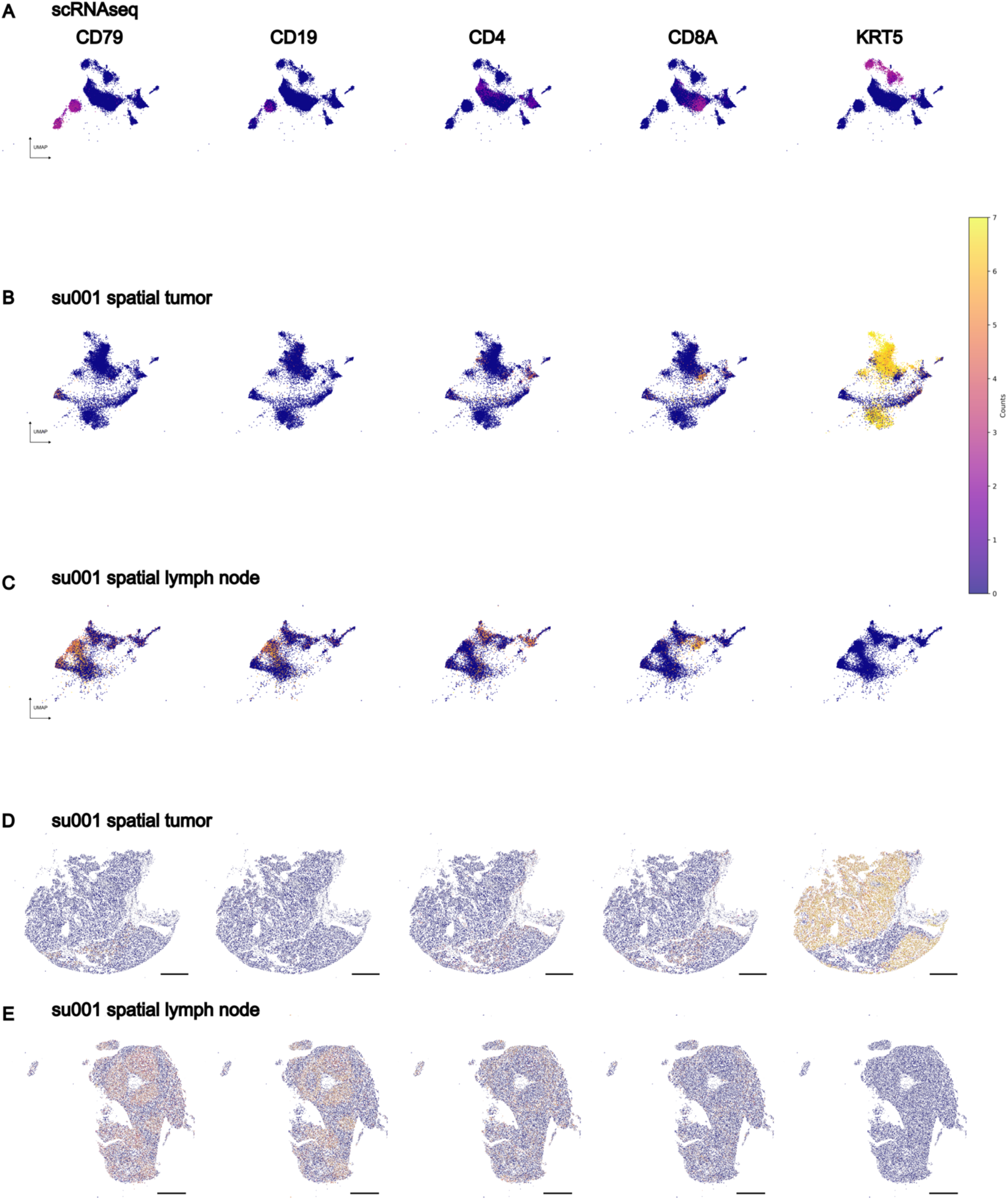
EVNI Latent Space and Spatial Representation of Example Tumor and Lymph Node. A) UMAPs of scRNA-seq cells B) Xenium tumor sample cells and C) Xenium draining lymph node cells colored by expression of select cell type markers in the shared ENVI latent embeddings. D) Spatial plots of the sample shown in B. E) Spatial plots of the sample shown in C. Scale bars are 0.5mm.

**Extended Data Figure 5:**
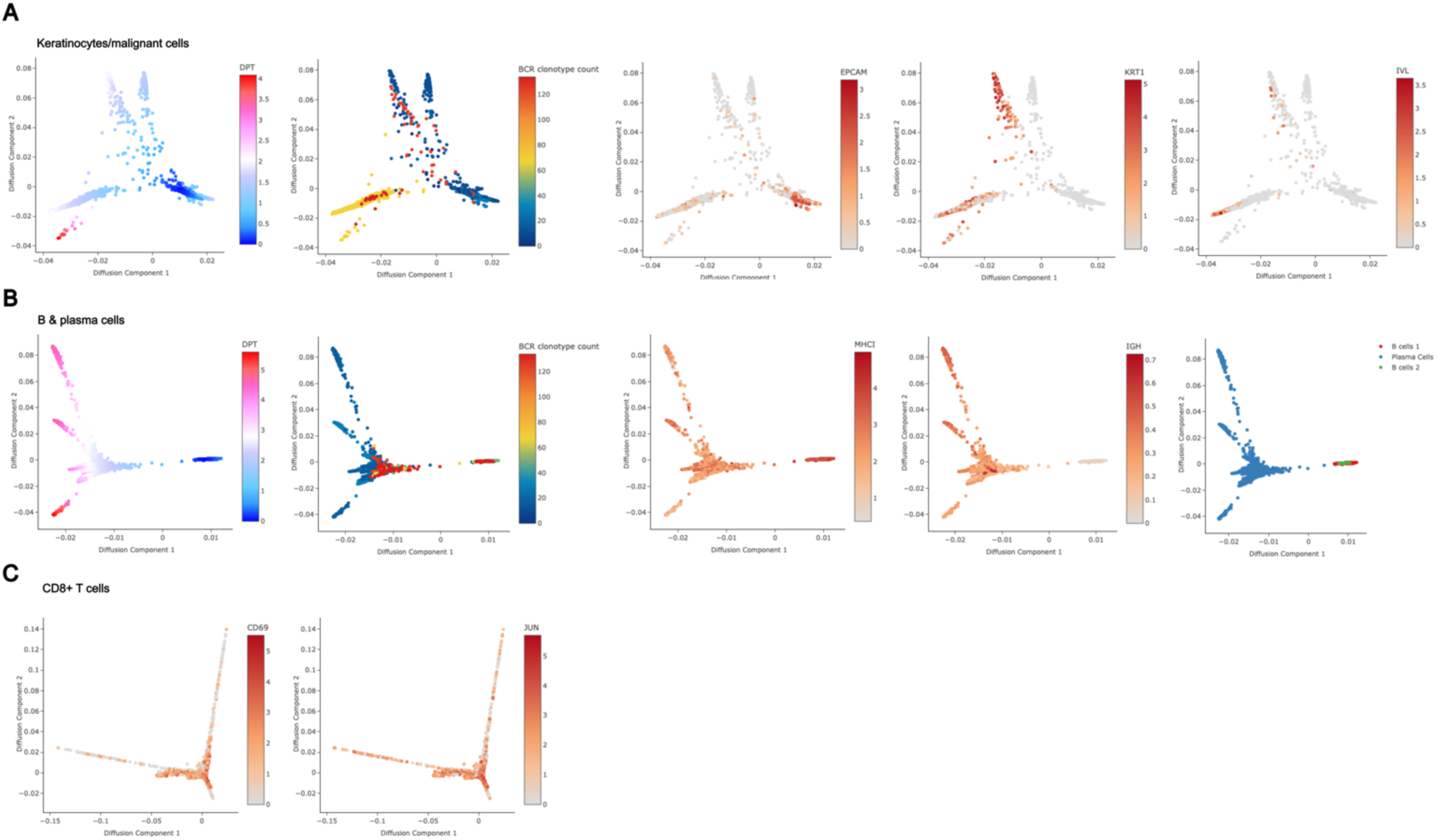
Diffusion Plots of Tumor and Immune Cell Clusters Reveals Increased Activated CD8+ T-cell representation and Decreased Tumor Burden in BCR-rich samples. A) Keratinocyte cells (identified in this study broadly as malignant cells) diffusion plot. DPT stands for diffusion pseudotime. The BCR clonotype count for each cell plotted is defined as the number of BCR clonotypes present in the tissue sample from which the plotted cell originated, representing the BCR diversity of its environment. Expression of EPCAM (BerEP4) a marker of basal cell carcinomas and progenitor keratinocytes, KRT1 an early differentiation marker, and IVL a late differentiation marker plotted per cell. B) B and plasma cells aggregated diffusion plot. MHC-I expression is defined as the mean expression of all class I MHC genes per cell. IGH expression is defined as the mean expression of all heavy chain Ig genes per cell. C) CD8+ T cells diffusion plot. Expression of CD69 and JUN, markers of TCR-induced activation plotted per cell.

**Extended Data Figure 6:**
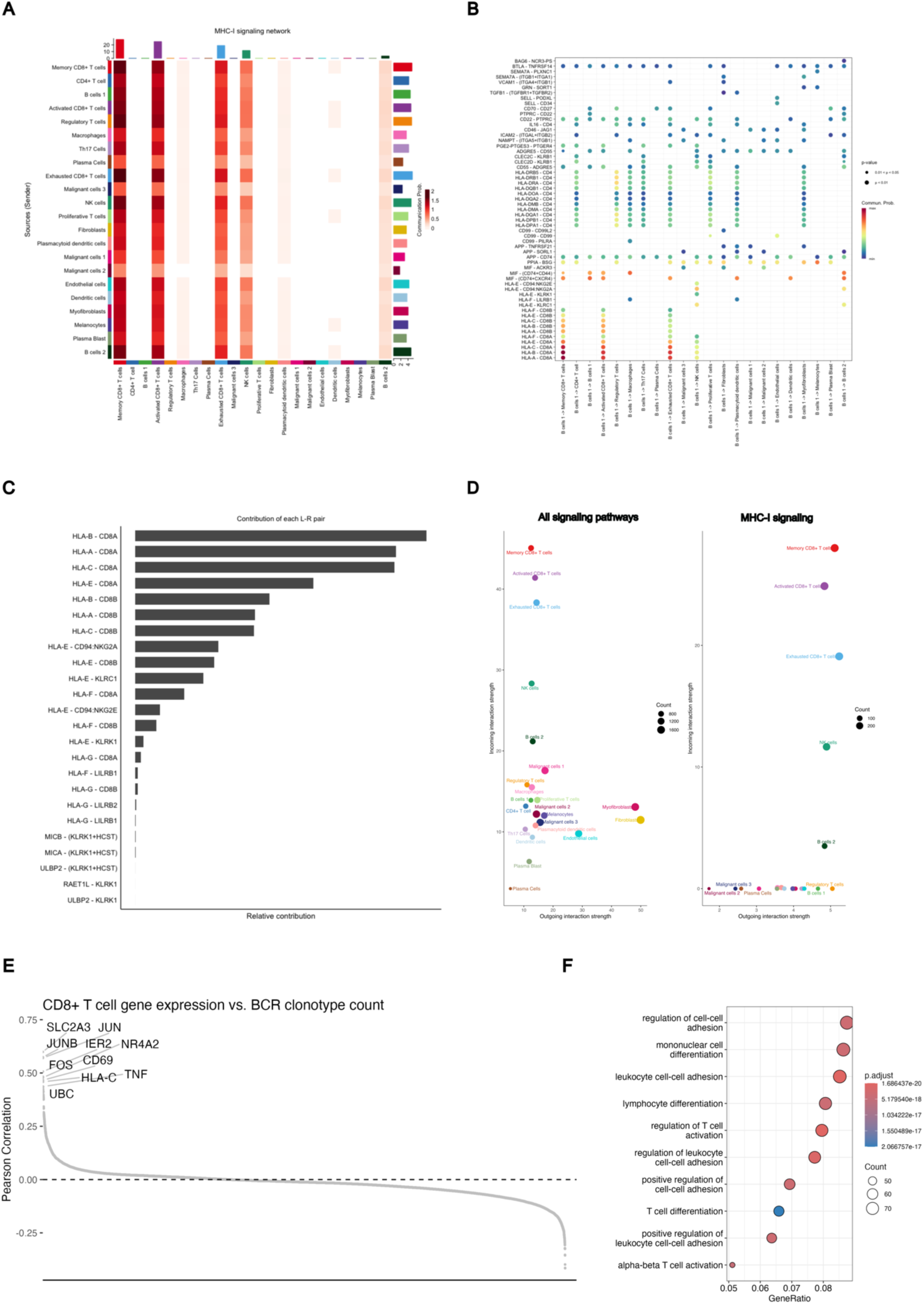
CellChat Analysis of B-Cell/T-Cell Signaling. A) Heatmap of communication probabilities in MHC-I signaling pathways between cell clusters. B) Significant communication pathways originating from cluster B cell 1 representative of B outgoing B cell signaling. C) Relative contribution of each ligand-receptor pair to MHC-I signaling strength between B cell clusters and Activated CD8+ T cells. D) Outgoing and incoming signaling strength per cell cluster for all signaling pathways (left) and MHC-I signaling pathways (right) calculated using CellchatDB. E) Pearson correlation between gene expression in CD8+ T cells vs. the BCR clonotype count in the respective tissue sample. Each point is the correlation of a single gene, and the plot it ordered from most positive correlation to most negative correlation. F) GO-Terms enriched in genes significantly correlated with BCR clonotype count in CD8+ T cells (defined as adjusted p-value <0.05).

**Extended Data Figure 7:**
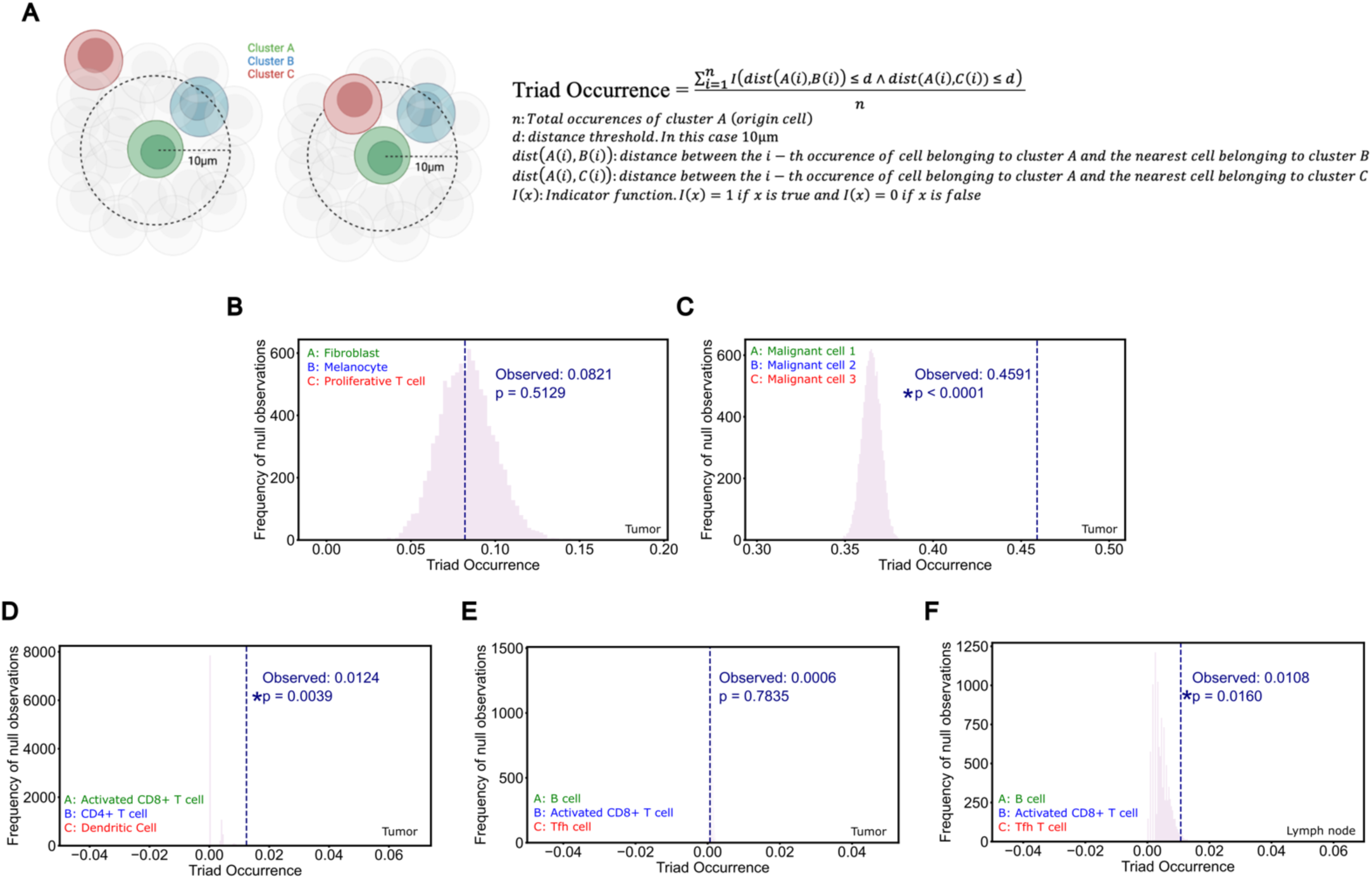
Immune Triads Involving B cells and Activated T cells. A) Schematic and equation of spatial cell triad calculation. The Triad Occurrence is defined as the proportion of existing co-localized pairs of two cell types (Cluster A and B) that also have a proximal third cell type interaction (Cluster C). B) Fibroblast/Melanocyte/Proliferative T cell Triad Occurrence; negative control in aPD1-treated tumors. C) Malignant cell 1/Malignant cell 2/Malignant cell 3 Triad Occurrence; positive control in aPD1-treated tumors. D) Activated CD8+ T cell/CD4+ T cell/Dendritic cell Triad Occurrence in aPD1-treated tumors. E) B cell/Activated CD8+ T cell/Tfh Triad Occurrence in aPD1-treated tumors. F) B cell/Activated CD8+ T cell/Tfh Triad Occurrence in aPD1-treated draining lymph nodes.

**Extended Data Figure 8:**
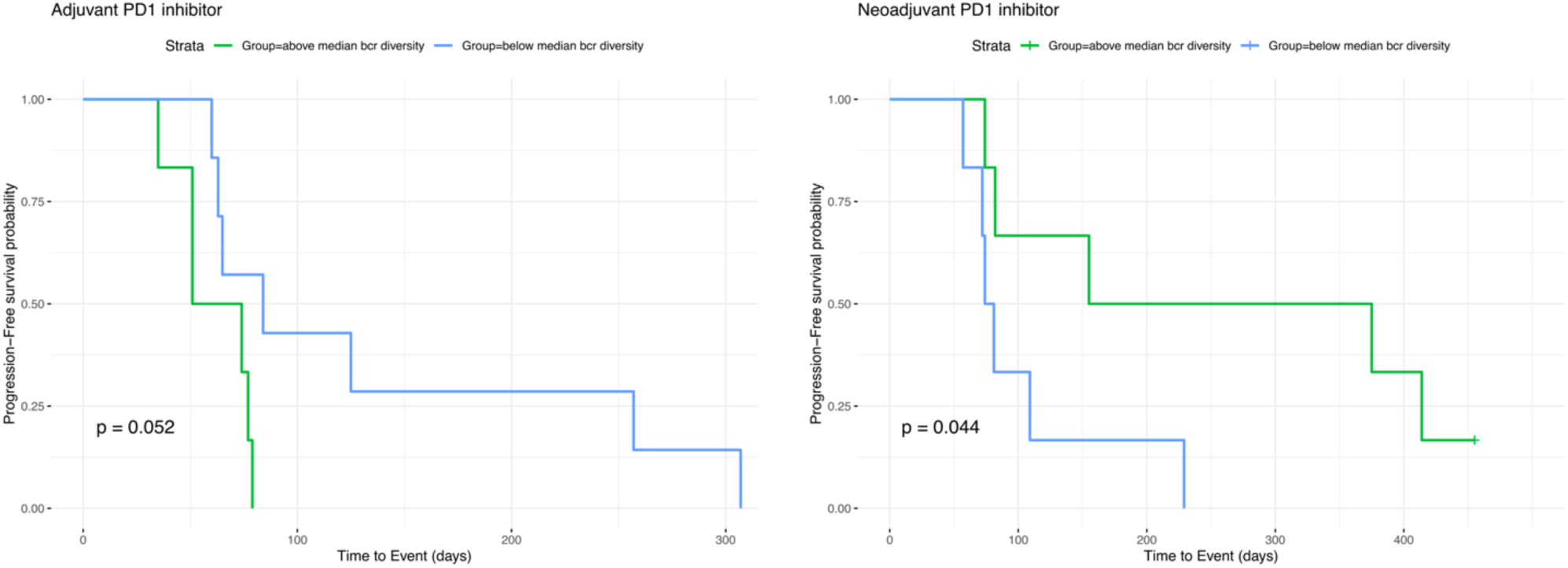
Progression-free Survival in Glioblastoma as a Function of PD-1 inhibitor-Induced BCR Diversity. Left: Progression-free survival in glioblastoma patients who underwent resection then started on adjuvant PD1 inhibitor. Tumor collection on which BCR clonotype analysis was performed was prior to PD1 exposure (Cloughesy et al. 2019). P values reported for log rank test. Right: Progression-free survival in glioblastoma patients exposed to neoadjuvant PD1 inhibitor followed by surgical resection.

